# Extended and dynamic linker histone-DNA interactions control chromatosome compaction

**DOI:** 10.1101/2020.10.10.334474

**Authors:** Sergei Rudnizky, Hadeel Khamis, Yuval Ginosar, Efrat Goren, Philippa Melamed, Ariel Kaplan

**Affiliations:** Faculty of Biology, Technion – Israel Institute of Technology, Haifa (32000), Israel; Faculty of Physics, Technion – Israel Institute of Technology, Haifa (32000), Israel; Russell Berrie Nanotechnology Institute, Technion – Israel Institute of Technology, Haifa (32000), Israel

## Abstract

Chromatosomes play a fundamental role in chromatin regulation, but a detailed understanding of their structure is lacking, partially due to their complex dynamics. Using single-molecule DNA unzipping with optical tweezers, we reveal that linker histone interactions with DNA are remarkably extended, with the C-terminal domain binding both DNA linkers as far as ~ ±140 bp from the dyad. In addition to a symmetrical compaction of the nucleosome core governed by globular domain contacts at the dyad, the C-terminal domain compacts the nucleosome’s entry and exit. These interactions are dynamic, exhibiting rapid binding and dissociation, sensitive to phosphorylation of a specific residue, and crucial to determining the symmetry of the chromatosome’s core. Extensive unzipping of the linker DNA, which mimics its invasion by motor proteins, shifts H1 into an asymmetric, off-dyad configuration and triggers nucleosome decompaction, highlighting the plasticity of the chromatosome structure and its potential regulatory role.

## Introduction

In eukaryotes, genomic DNA is packaged into nucleosomes, comprised of ~147 base pairs (bp) of DNA wrapped ~1.65 times around an octamer of the core histone proteins H3, H4, H2A, and H2B^1–3^, and arranged as ‘beads on a string’ separated by linker DNA^4,5^. Further organization is supported by one of several linker histones (H1s)^6^, which bind the nucleosome to form a chromatosome^7^, facilitating the formation of higher-order structures^8,9^ and promoting liquid-liquid phase separation^10^. Despite the central role that linker histones play in organizing and regulating the chromatin structure, a molecular understanding of their function is only starting to emerge^11–14^.

Eleven subtypes of linker histones have been identified in humans and mice, and they share a conserved structure, composed of a winged-helix globular domain (GD) flanked by unstructured and intrinsically disordered short N-terminal (NTD) and long C-terminal (CTD) domains^12,15^. The exact position and nature of the interactions between H1 and the nucleosome remain controversial^16–21^, and their precise contribution to chromatin compaction is a matter of debate^22–25^. Available crystal structures show two binding modes for the GD, which differ in their symmetry relative to the dyad (the center of the nucleosomal DNA). In the “on-dyad” mode, the GD binds the nucleosome at three distinct DNA binding sites, one at the center, and the other two at the entry and exit positions^24,26,27^. In the “off-dyad” mode, the GD interacts with the nucleosome at two primary locations, ~10 bp off-dyad and with a single DNA linker^28,29^. A previous study showed that the orientation of H1 is dictated by the amino acid sequence specific for a subtype^30^; however, both binding modes were observed for the same variant using different experimental approaches^16–21^. Even more intriguing is the nature of the CTD interaction with the nucleosomal DNA. Earlier studies indicated that although the CTD is intrinsically disordered, it becomes partially ordered when engaged with the DNA^31–33^. In contrast, more recent studies show that the CTD remains disordered on a nucleosome^34–36^. Moreover, while the position of the CTD mapped to both DNA linkers by hydroxyl-radical footprinting^17^, a recent cryo-EM study showed that it is associated with only a single linker^26^.

The exterior positioning of H1 on the nucleosome makes it highly mobile^37,38^, promoting structural plasticity of chromatin fibers^26,27,29^. At the level of a single-nucleosome, H1 suppresses the spontaneous wrapping and unwrapping (“breathing fluctuations”) of linker DNA^26^ and accelerates the folding of the outer wrap^39^. However, H1 can also spontaneously detach from the linker DNA^26,40^, leading to increased breathing of the entire particle. These transitions were recently shown to play a critical role in transcription regulation^39,40^; however, their dynamic nature presents a challenge for studying the chromatosome structure using traditional biochemical approaches, with their inherent ensemble averaging. Moreover, high-resolution structural studies often require truncation of H1 and shortening of the linker DNA, or even cross-linking, thus obscuring the underlying and functionally important dynamics.

In a previous work, Wang and colleagues used a single-molecule approach based on DNA unzipping with optical tweezers to map the strength of histone-DNA interactions inside the nucleosome^41^. This approach was later used to study the properties of centromeric nucleosomes^42^ and nucleosomes containing the variant H2A.Z^43^, and how the structure of the nucleosome affects the crossing dynamics of Pol II^44^. We further exploited this idea and developed an assay to measure the dynamic repositioning of a nucleosome by ‘partial’ DNA unzipping^43,45^. Unfortunately, the application of these approaches to chromatosomes is challenged by the spontaneous dissociation of linker histones under single-molecule conditions^46^. In our current work, we bypassed this problem by reconstituting chromatosomes *“in situ”,* under singlemolecule conditions, using a laminar flow cell^47,48^. Subjecting chromatosomes to DNA unzipping revealed that the stabilization and symmetry of their structure are supported by extended and dynamic interactions. Although a symmetric and compact on-dyad structure is the more stable one, other configurations form upon perturbation of these interactions, such as phosphorylation of specific residues or invasion by cellular machinery. Together, these results shed light on the contribution of different structural elements to the dynamic control of DNA packaging and highlight a potential regulatory role for the chromatosome.

## Results

### DNA unzipping reveals nucleosome compaction by linker histone

To study the effect of linker histone on nucleosome compaction, we reconstituted nucleosomes on the Widom 601 positioning sequence^49^ and ligated them to terminally modified DNA handles connected to two beads trapped by high-resolution optical tweezers^43,47^. After tethering a single nucleosome between the trapped beads, we exploited a laminar flow cell to move the construct to a region containing 5 nM of full-length H1^0^ (from now on referred to as H1) to form a chromatosome *in situ.*

Subsequently, we moved the chromatosomes to an H1-free channel to avoid multiple binding events or formation of non-native aggregates, and immediately subjected them to full and irreversible unzipping to prevent spontaneous dissociation of H1 (Fig. 1a). Propagation of the unzipping fork led to the sequential disruption of protein-DNA interactions, revealing their position and strength, starting with those associated with the linker DNA, which are not in direct contact with core histones (Fig. 1b). Next, the interaction of the N-terminal part of H3 (H3-NTD), located ~-80/-70 bp away from the dyad, at the nucleosome’s entry, is disrupted. Since this interaction is highly dynamic, it is generally undetected when unzipping the nucleosome. Remarkably, following exposure to H1, the interaction was clearly detected, and the force required to disrupt this interaction was increased (Fig. 1c), indicating stabilization of this contact, consistent with the reported reduction of DNA ‘breathing’^26^. Unzipping the DNA further led to the rupture of the strong interactions between the DNA and the proximal H2A/H2B dimer located ~-60/-40 bp off-dyad, and the strongest interactions with the (H3/H4)2 tetramer positioned −20 next to the dyad^41^, indicating that the presence of linker histone does not alter the position of primary histone-DNA contacts. However, the force required to overcome these interactions was elevated, suggesting that H1 compacts the DNA also at the nucleosomes’ core (Fig. 1c,d). To explore this stabilization effect in a biologically relevant DNA sequence, we assembled nucleosomes using a DNA fragment of the *Cga* gene, previously shown to harbor a nucleosome *in vivo*^43^. When exposed to H1, we observed a similar stabilization as that observed with 601 nucleosomes (Fig. 1e), both at the entry and the core, suggesting that H1 compacts similarly the nucleosomes formed on naturally occurring DNA. Interestingly, we noticed that our ability to detect the relatively weak interactions at the linker DNA was significantly improved for *Cga* relative to 601 nucleosomes. When we analyzed the traces, it was evident that the ‘background’ unzipping force of naked DNA was significantly reduced for *Cga* (Fig. S1a), likely due to its lower GC content. Hence, in order to combine the precise nucleosome positioning provided by the 601 sequence with a reduced background that allows detection of weaker interactions at the linker DNA, we designed a new DNA construct (which we termed 601-AT) composed of the central 73 bp of 601 DNA, responsible for its positioning properties^50^, flanked by two identical ~184 bp fragments of AT-rich DNA (Fig. S1b, inset). A thermodynamic model of the unzipping reaction, based on the base-pairing energy and the DNA flexibility^51^, predicts a substantial drop in the unzipping force within the altered regions (Fig. S1b). Indeed, when we unzipped the modified DNA construct we observed a ~ 6 pN decrease in the rupture force at the corresponding DNA regions (Fig. S1a,c). Next, we performed several control experiments to ensure that our modulation didn’t affect the nucleosome’s position and structure. 601-AT nucleosomes showed a single population in a gel-shift assay (Fig. S1d) and positional dispersion comparable to 601 nucleosomes under single-molecule conditions (Fig. S1e). Moreover, their unzipping signature showed the two known regions of strong interaction at the expected locations^41,45,52^, reflecting the particles’ structural integrity (Fig. S1c). Interestingly, the weak interaction of H3-NTD that was generally masked in 601 nucleosomes appeared to be more pronounced, consistent with the increased sensitivity of the 601-AT construct (Fig. S1c). Finally, nucleosomes formed on 601-AT DNA were able to produce chromatosomes both in bulk and singlemolecule conditions in situ, when we prepared them using commercially available human H1, or its bacterially expressed mouse paralog (Fig. S1f-i).

**Figure 1.**
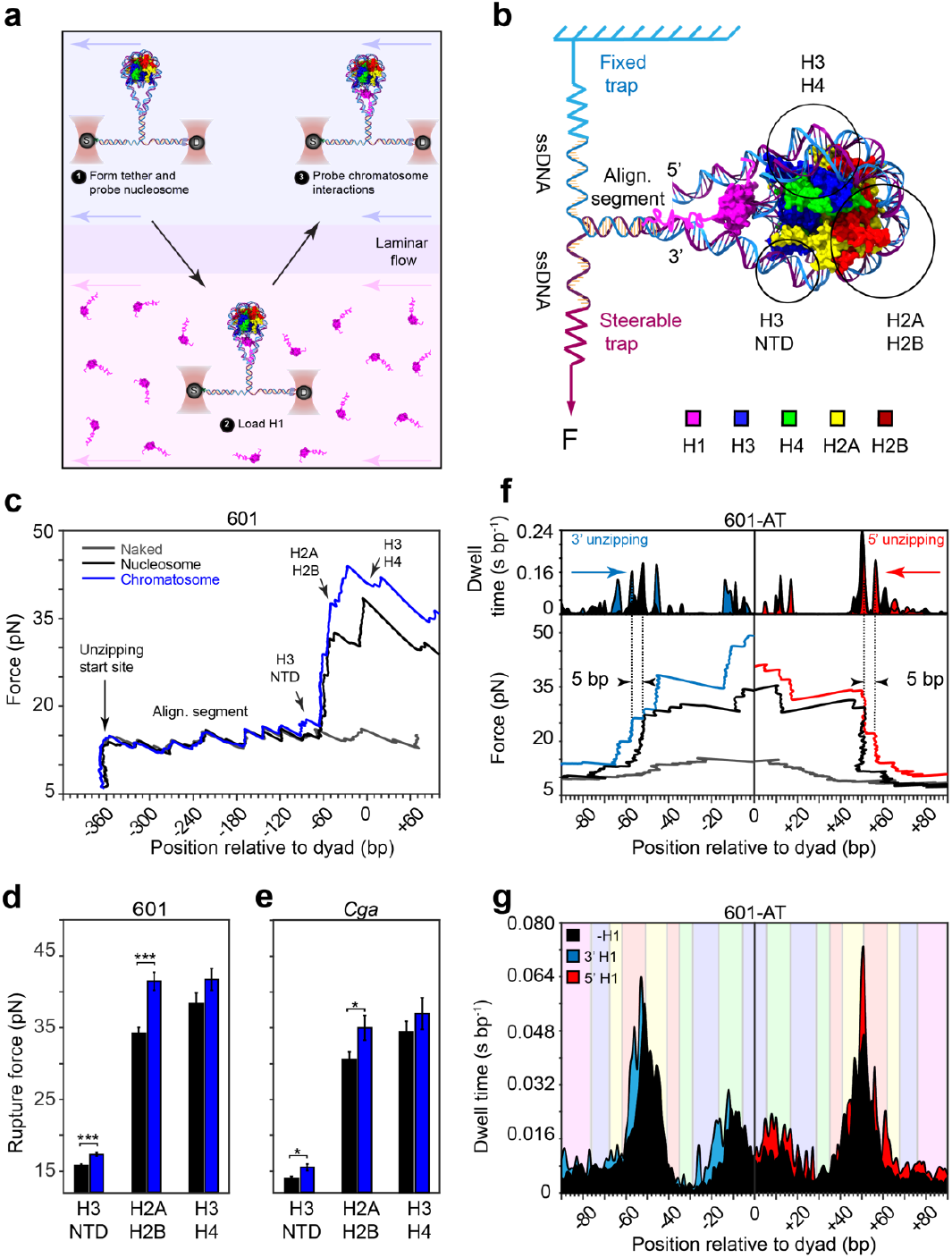
DNA unzipping reveals nucleosome compaction by linker histone. a) Schematic description of the experimental assay. The nucleosome is tethered between Streptavidin (S), and anti-Digoxigenin (D) coated beads captured by dual-trap optical tweezers. The trapped nucleosome is exposed to [5 nM] H1 to form a chromatosome and then subjected to DNA unzipping in the H1-free channel. b) Representation of the DNA unzipping reaction through a chromatosome, based on the crystal structure of the GD bound to a 197 bp palindromic 601L nucleosome (PDB: 5NL0)^26^. Hypothetical positions for H1 CTD and NTD domains are shown for clarity. Two strands of the DNA are connected to DNA handles bound to the trapped S and D beads. Moving one trapped bead away from the other creates tension, leading to the conversion of dsDNA to ssDNA, which allows probing the position and strength of major histone-DNA interactions (circled). c) Representative unzipping curves for nucleosomes reconstituted using the 601 DNA without (black) or in the presence of H1 (blue). ‘Naked’ (i.e. no nucleosome or H1) 601 DNA (grey) is shown for reference. The unzipping reaction starts at ~-360 bp from the dyad, proceeds through the fixed ‘alignment segment’, reaching histone-DNA interactions in a chromatosome as highlighted in Fig.1b. d,e) Mean rupture forces for H3-NTD, H2A/H2B, and H3/H4 interactions, shown for 601 (d) and Cga (e) nucleosomes. Data shown as mean±s.e.m.; n_601=13_, n_601+H1_=15, n_*Cga*_=14, n_*Cga*+H1_=12. \**P*<0.05, \*\*\**P*<0.001, two-sample Student’s *t*-test. f) Nucleosomes formed on the modified 601-AT construct were unzipped from the 3’ or 5’ end in the absence (black) or presence of H1 (blue or red, respectfully). Naked DNA is shown for reference (grey). Single representative traces for each condition (bottom panel) with their corresponding dwell time histograms (top panel) are shown for 3’ unzipping (left) and 5’ unzipping (right). Note a ~ 5 bp periodicity pattern of interaction in the nucleosome that is also conserved in the presence of H1. g) Average dwell time histograms constructed from multiple traces (n_5’,-H1_=46, n_5’,+H1_=21, n_3’,-H1_=53, n_3’,+H1_=41). Shaded colors (according to the color code in Fig.1b) indicate positions of interactions of each histone as inferred from the crystal structure.

When chromatosomes were formed using the 601-AT construct in situ, we observed a stabilization similar to that observed for the 601 nucleosomes, but with higher sensitivity, as reflected by a ~5 bp periodicity of interactions, which were clearly stabilized in the presence of H1 (Fig 1f, left-bottom panel). Conversion of the individual traces’ force-position data to dwell times (Fig. 1f, left-top panel), and averaging over all the traces in the data set (Fig. 1g, left panel), allowed us to generate a ‘compaction map’ of nucleosomal interactions stabilized by H1 (Fig. 2a). Notably, although the 5 bp periodic pattern within the interaction cluster was similar to that of nucleosomes, the magnitude of dwell times was significantly higher for chromatosomes, particularly −50 bp and −20 bp away from the dyad. Notably, these experiments probe one side of the nucleosome, which appears to be stabilized by H1. This is consistent with H1 binding the nucleosome asymmetrically, in the ‘off-dyad’ conformation, as proposed by several previous studies^17–19,22,25,26^. However, when we unzipped the complexes from the opposite (5’) end, we observed a similar stabilization of the H3-NTD contact as with the 3’ end unzipping experiments (Fig. 1f,g, right panel; Fig. 2a), indicating that H1 contacts the DNA on both sides of the nucleosome. Moreover, the H3/H4 and H2A/H2B regions were stabilized to a similar extent in the 3’ side, suggesting that H1 binds the nucleosome in an on-dyad configuration, compacting the entire structure symmetrically.

**Figure 2.**
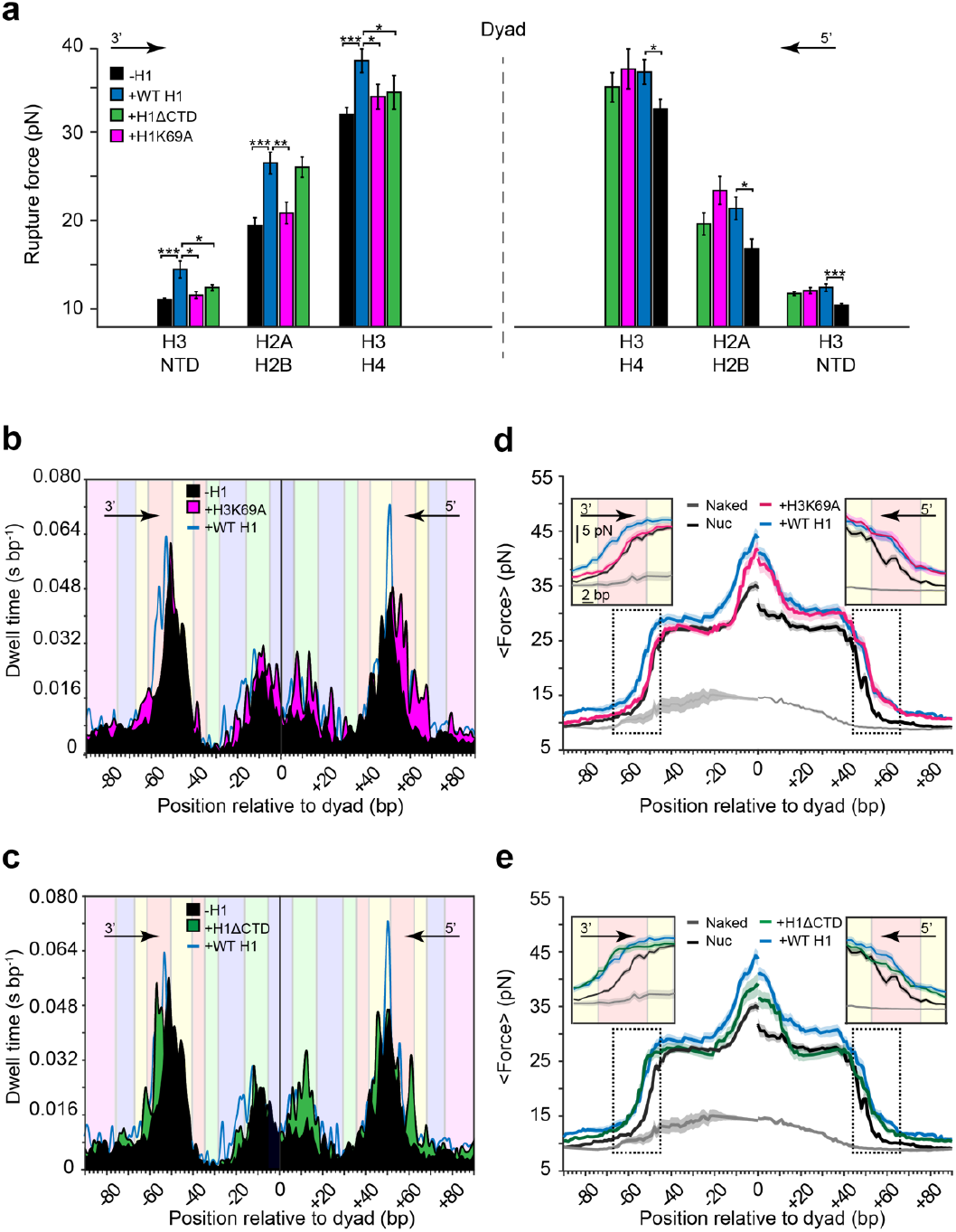
H1-DNA contacts at the dyad are responsible for symmetric compaction. Unzipping traces obtained from 3’ and 5’ unzipping experiments with the H1K69A (magenta) and H1ΔCTD (green) mutants were analyzed and are presented as a) mean rupture force of regions of interactions, b-c) dwell time histograms and d-e) average force-position curves. Dwell time histograms of nucleosomes (black) and chromatosomes formed with WT H1 (blue) are identical to those in Fig.1g and shown for reference. n_5’-H1_=46, n_5’nuc+H1_=21, n_3’-H1_=53, n_3’nuc+H1_=41, n_5’nuc+H1K69A_=22, n_3’nuc+H1K69A_ = 27, n_5’nuc+H1ΔCTD_=22, n_3’nuc+H1ΔCTD_ = 20. \**P*<0.05, \*\**P*<0.01, \*\*\**P*<0.00, two-sample Student’s *t*-test.

### H1-DNA contacts at the dyad are responsible for symmetric compaction

To understand what region in H1 is responsible for the symmetric compaction, we decided to focus on the dyad-binding residues. Among them is a conserved lysine at position 69, which interacts with one of the seven bp at the nucleosome center^26^, and is crucial for H1 binding *in vitro*^24^ and *in vivo*^38^. To test whether this contact is involved in the observed stabilization, we mutated it to alanine and expressed the altered H1 (H1K69A) in *E. coli* (Fig. S1f). We then used the purified variant to form chromatosomes *in situ,* followed by DNA unzipping from the 5’ or 3’ orientation. When probed in the 5’ direction, binding of the mutated H1 resulted in a similar stabilization of the nucleosome as with WT H1, as indicated by higher forces and elevated dwell times within the H2A/H2B and H3/H4 regions (Fig. 2a,b, right panels). However, when we examined chromatosomes from the 3’ end, the stabilization of the dyad interaction was significantly reduced (Fig. 2a,b, left panel), not only for the H3/H4 region proximal to the mutated lysine 69, but also for the relatively distant H2A/H2B contacts. This asymmetry between 3’ and 5’ halves was clearly seen also in average force-position curves (Fig. 2d). Strikingly, even the more distant H3-NTD-DNA interaction located ~80 bp away from the mutated position, was reduced in the 3’ side (Fig. 2a,b,d), suggesting that the residues at the dyad are essential for stabilizing the symmetric ‘on-dyad’ configuration. This result is consistent with a previous study, which showed that residues close to the dyad could determine whether the chromatosome is in the on- or off-dyad modes^30^.

Conversely, previous studies showed that truncation of the CTD does not affect the binding orientation^24,26,30^. Hence, to further explore the contribution of the H1 CTD to the observed stabilization, we expressed and purified the 1-97 amino acid fragment of H1 lacking the CTD (H1 ΔCTD, Fig. S1f) and repeated the unzipping experiments with the truncated variant. Although we observed a clear destabilization at the entry, and a subtle one at the core and exit regions, H2A/H2B contacts remained stabilized symmetrically around the dyad, similarly to when chromatosomes contained the WT H1 (Fig. 2a,c,e). This implies that although the CTD encompasses almost half of H1, it does not affect the symmetry of compaction, but instead supports the existing H1-DNA contacts.

### Dynamic mapping uncovers extended interactions of H1 CTD and linker DNA

The weakening of chromatosome compaction upon the deletion of the CTD motivated us to explore further the nature of the H1 CTD interaction with a nucleosome. We decided to focus on contacts with the linker DNA, where the CTD was proposed to bind^17,26^. Although the presence of interactions in this region was evident from the initial experiments, their relatively weak rupture forces and likely rapid dynamics prevented us from systematically studying them using an irreversible unzipping experiment. Hence, we developed an assay based on partial unzipping, which probes the position and strength of H1-linker DNA interaction multiple times in the same molecule. Of note, repetitive partial unzipping does not destabilize the nucleosome or induce repositioning, as shown in our previous work^45^. Nucleosomal DNA was unzipped until reaching, but not disrupting, the proximal H2A/H2B interaction (typically ~19 pN). After the interaction was detected, the force was relaxed again, and the construct rezipped (Fig. 3a). Next, the same nucleosome complex was exposed to H1 to form a chromatosome, moved to a region depleted of H1, and subjected to ~10 cycles of reversible unzipping (~8 s for each cycle). Data collection efficiency using multiple unzipping cycles was now significantly increased. Furthermore, our data analysis was significantly improved by using control experiments with the same nucleosome for alignment.

**Figure 3.**
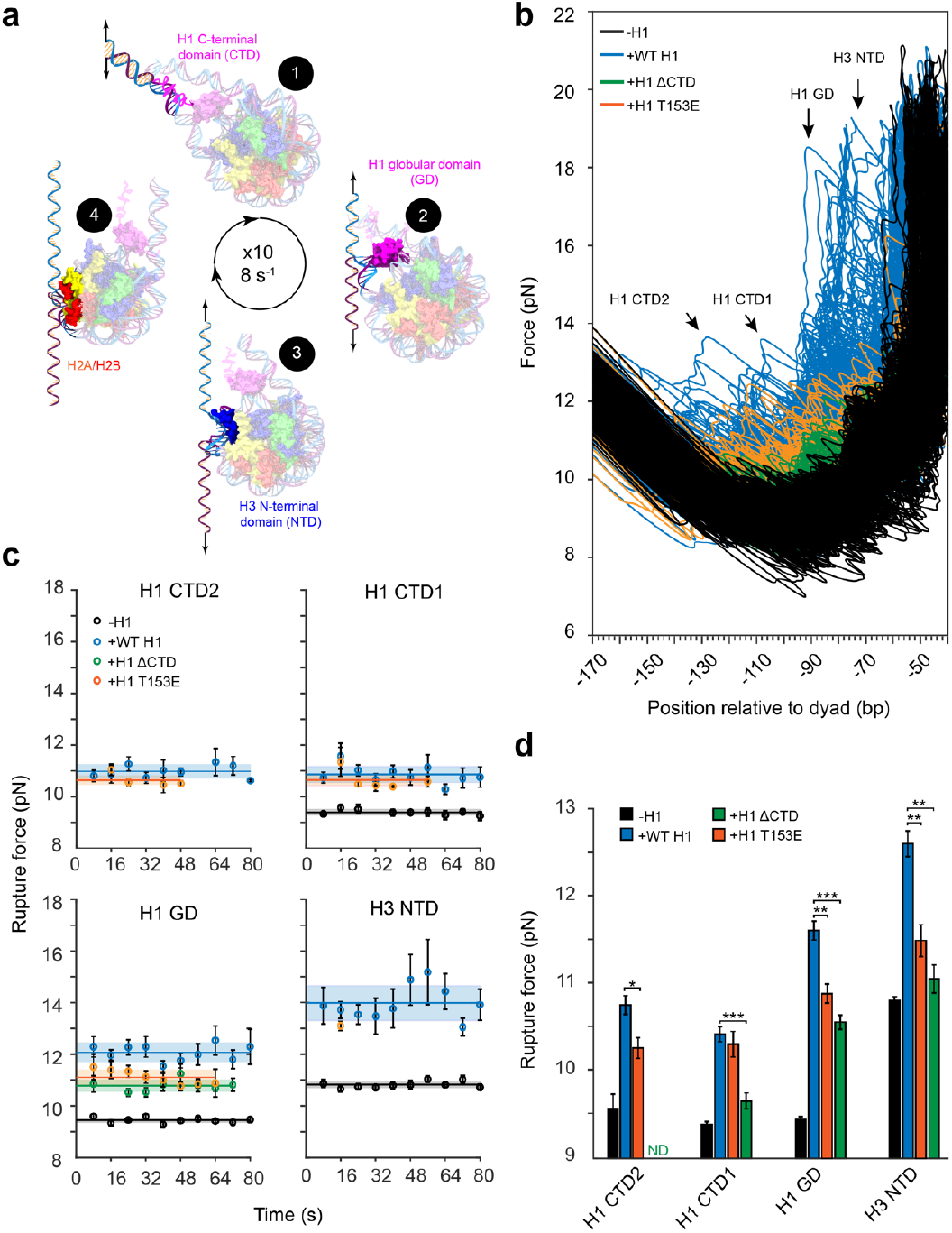
Dynamic mapping uncovers extended interactions of H1 CTD and their role in nucleosome compaction. a) Schematic description of the reversible unzipping assay to probe H1 interactions with linker DNA. Progression of the unzipping fork displaces H1 directionally, starting with (1) the CTD, followed by (2) the GD and (3) H3-NTD, until reaching an H2A/H2B dimer (4). Next, the tension is relaxed and dsDNA is reformed, after which the unzipping cycle is repeated (10 times, 8 s per cycle). b) Repetitive unzipping force-extension curves of the WT (blue), H1ΔCTD (green) and H1T153E(orange) chromatosomes or nucleosomes without H1 (black), all unzipped from the 3’ direction. The WT chromatosomes were probed under STO conditions. The H1ΔCTD and H1T153E binding to nucleosomes was unstable overtime (Supp. Fig. 2c) and hence investigated using MTO conditions. The clusters of detected interactions are highlighted with arrows. The interactions measured in each unzipping cycle (see Methods) were quantified and presented as c) mean rupture forces as a function of time within an 8 s window, and d) mean rupture forces. Data are shown as mean±s.e.m.; n_nuc_=307, n_nuc+WT H1_=252, n_nuc+H1ΔCTD_=41, n_nuc+H1T153E_=54. \**P*<0.05, \*\**P*<0.01,\*\*\**P*<0.001, two-sample Student’s *t*-test.

Unzipping curves now clearly showed the presence of force rips in the linker DNA region, which can be clustered into four defined areas of interactions (Fig. 3b). The mean rupture force for each interaction remained constant over time (Fig. 3c), suggesting that they can rapidly regain their native conformation upon relaxation and rezipping. The position of the H3-NTD contact at the entry, ~-70 bp away from the dyad^3^, was readily detected also without H1 (Fig. 3b), albeit with significantly lower forces (Fig. 3c,d), consistent with the quantified rupture forces measured in the previous irreversible unzipping experiments (Fig. 1d,e; Fig. 2a). Three additional interaction regions upstream did not appear in the absence of H1. At approximately −90 bp, we detected an interaction region that, based on the available crystal structures^26,28^, we attribute to the GD-linker DNA interaction. Interestingly, approximately −115 bp and −140 bp away from the dyad, we observed two previously unreported interactions, which we suspected were formed by CTD (CTD1 and CTD2; Fig. 3b, Fig. 4a). Indeed, these contacts were absent in ΔCTD chromatosomes (Fig. 3b,d), confirming that they belong to the CTD. The mean rupture force of these interactions was significantly lower than for the GD, suggesting that the CTD is more loosely bound to the linker DNA. Similar clusters of interactions were also observed when we performed the experiments with *Cga* nucleosomes, but were less pronounced for 601 ones, likely due to the higher GC content of 601 DNA (Fig S2a,b). Notably, these newly detected contacts between H1 and linker DNA indicate that the size of the chromatosome is significantly larger than previously reported.

**Figure 4.**
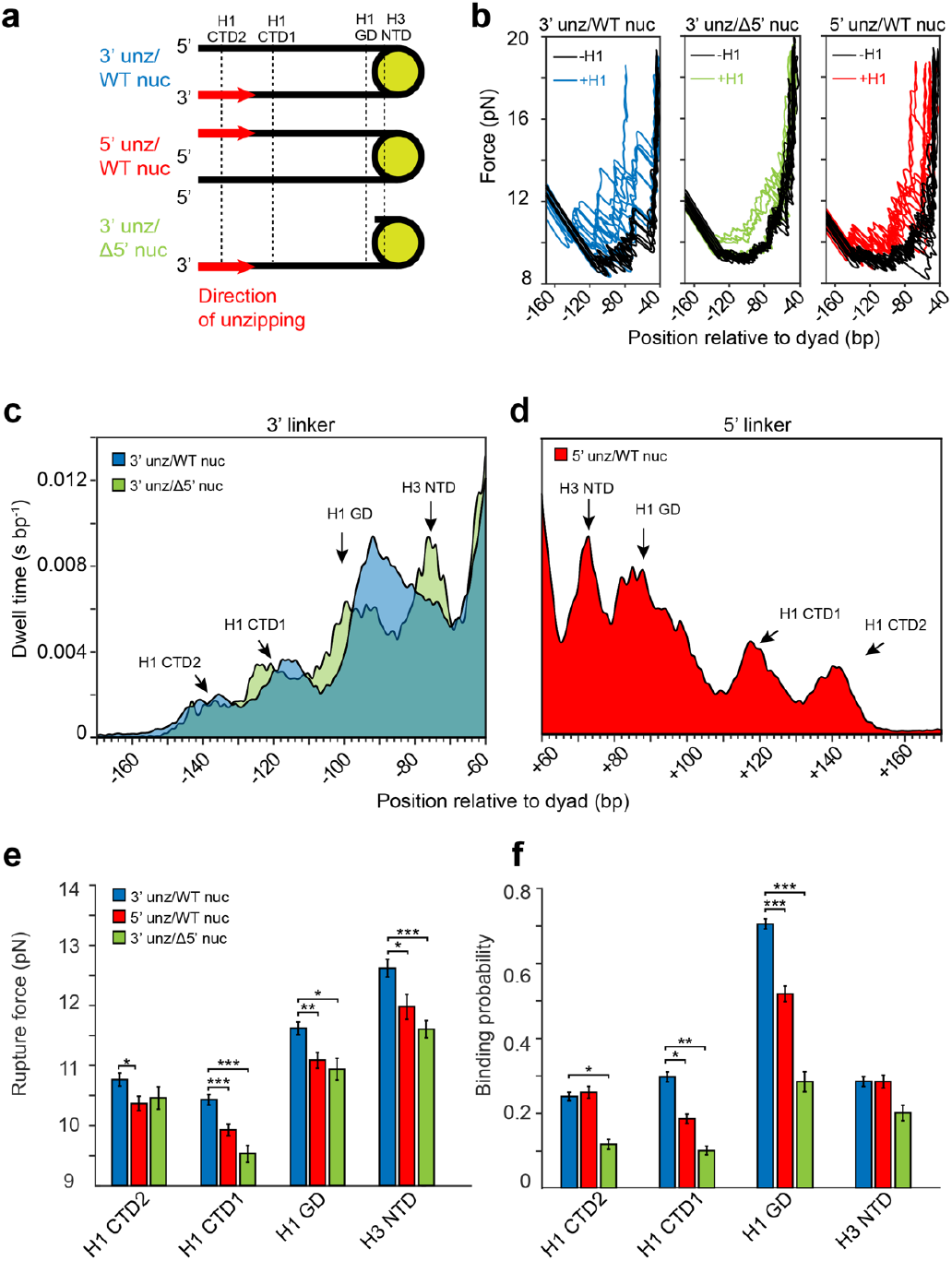
H1 dynamically interacts with both linkers to stabilize the on-dyad conformation. a) Molecular constructs used for repetitive unzipping experiments. 3’ unz/WT nuc – WT nucleosome unzipped from 3’ end. 5’ unz/WT nuc – WT nucleosome unzipped from 5’ end. 3’ unz/Δ5’ nuc - nucleosome harboring a full 3’ linker to be unzipped, but with a 3 bp long 5’ linker, designed to abolish interactions with GD, CTD1 and CTD2. b) Representative traces of WT or single-linker chromatosomes unzipped partially from the 3’ or 5’ direction. Corresponding control experiments without H1 are shown in black. c-d) Average dwell time histograms constructed from partially unzipped traces (~10 cycles, 8 s each) for Δ5’ chromatosomes (c, green) and WT chromatosomes probed from the 3’ (c, blue) and 5’ (d, red) direction. All traces were low-passed filtered to 150 Hz and binned to 1 bp. The clusters of detected interactions are highlighted with arrows. e) Rupture forces and f) binding probabilities calculated for the clustered interactions, as shown in Fig. 4 c,d. Data are shown as mean±s.e.m.; n_5’nuc_=136, n_3’nuc_=187, n_Δ5’nuc_=60, n_5’nuc+H1_=139, n_3’nuc+H1_=242, n_Δ5’ nuc+H1_=60. \**P*<0.05, \*\**P*<0.01,\*\*\**P*<0.001; two-sample Student’s *t*-test for fig. 4e, χ^2^ test for Fig. 4f.

### CTD anchors the globular domain and stabilizes the nucleosome entry

Next, we aimed to explore the role of the detected CTD interactions in nucleosome compaction. However, repeating the reversible unzipping experiment with the ΔCTD mutant led to a rapid decrease in the fraction of successive binding events (i.e. the “binding probability”) over time (Fig. S2c), suggesting that its faster dissociation prevented us from characterizing its interactions under these ‘single-turnover’ (STO) conditions. To overcome this, we incubated the nucleosome with the mutant H1 under ‘multiple-turnover’ (MTO) conditions, with continuous exposure of H1, to form the complexes simultaneously. A lack of a decrease in binding probability in the GD position over time indicated the rapid formation of the complexes (Fig. S2c). In contrast, the rupture forces were similar between the MTO and STO (Fig. S2d), indicating that similar complexes form. Notably, in addition to eliminating the stabilization of the CTD interactions, both GD and H3-NTD interactions were significantly destabilized for the ΔCTD mutant (Fig. 3d).

Having established the importance of the entire CTD, we sought to explore the contribution of individual CTD-DNA contacts. Several reports indicate that the CTD binds the linker DNA via specific and structured S/TPKK domains, rather than contacting it non-specifically^31–33^. Mutation of threonine to glutamic acid, which mimics phosphorylation in this residue, reduces H1.1 binding *in vivo,* reminiscent of the effect of partial truncation of the CTD^53^. Accordingly, we decided to substitute the conserved threonine at position 153, located at the TPKK domain in H1, with glutamic acid (H1 T153E). We expressed and purified the mutant protein, assembled the chromatosome in situ, and subjected it to partial unzipping analysis under MTO conditions to prevent any possible dissociation. The rupture forces associated with CTD2, and the distant GD and H3-NTD (but not CTD1) were reduced, suggesting that phosphorylation induces a global conformational change (Fig. 3c,d). Overall, these results indicate that CTD allows H1 to stay on the DNA for a prolonged period of time and that, once engaged, mediates the GD compaction of the nucleosome at the linker region.

### H1 dynamically interacts with both linkers to stabilize the on-dyad conformation

The increase in compaction at both entry and exit sites (Fig. 2a) suggests that H1 also interacts with the opposite, 5’ linker DNA. To map interactions in this region, we conducted a partial unzipping experiment from the opposite orientation (5’unz/WT nuc - Fig. 4a). Contacts attributed to the GD were detected ~+90 away from the dyad (Fig. 4b, right panel), similar to unzipping the 3’ linker and consistent with the irreversible unzipping experiments. Interestingly, similarly to the proximal linker DNA, we detected clusters of interactions located +115 and +140 bp away from the dyad, suggesting that the CTD interacts with both linker arms at similar positions (Fig. 4c,d). Quantification of the mean breaking forces of the interactions, revealed slightly reduced values for both the GD and CTD in the 5’ orientation, suggesting a weaker binding to DNA (Fig. 4e).

The presence of interaction clusters at similar positions at both linkers could reflect two extreme scenarios or a combination of them. One possibility is that H1 binds both linkers similarly and concurrently, creating a ‘stem-like’ structure as proposed previously^17^. An alternative option is that H1 dynamically switches position between the two linkers, interacting with a single-linker at a time. To clarify this, we used the data from the reversible unzipping experiments to calculate the binding probability of each H1 interaction with the 5’ and 3’ linker. Notably, these probabilities, for all the interaction regions, are independent of the “incubation time”, i.e. the time before the interaction is probed (Fig. S3a), indicating that the system is in thermodynamic equilibrium and the measured probability reflects the energy of the interaction^48,54^. All interactions showed binding probabilities lower than 1, suggesting that H1 occasionally detaches from the linker DNA. If H1 were to alternate between the proximal and distal linker DNA, we would expect the sum of the probabilities to detect a given interaction, at the 5’ and 3’ side, to be also smaller than 1. However, the binding probability of the GD contact was ~0.7 for the 3’ linker and ~0.5 for the 5’ linker (Fig. 4f), suggesting that although the GD binds more weakly to the latter, for a significant fraction of time the GD interacts with both linkers simultaneously. The binding probabilities of CTD1 and CTD2 with the 3’ and 5’ linkers were substantially lower than for the GD (Fig. 4f), consistent with their dynamic nature.

To explore further the nature of GD and CTD contacts with DNA linkers, we decided to investigate H1 binding to nucleosomes lacking a 5’ linker (3’unz/Δ5’ nuc - Fig. 4a, Fig. S5a). In the case of simultaneous binding to two linkers, the deletion of one linker is expected to weaken the interactions with the remaining one, resulting in a *decrease* in their binding probability. In contrast, if H1 occupies only a single linker at a given time, we anticipate an *increase* in binding probability, since now the “competitor” linker is eliminated. When we partially unzipped the deletion nucleosomes in the presence of WT H1, an apparent elevation of force in the region corresponding to GD-DNA interaction was readily detected (Fig. 4b,c,e), confirming that it was able to form a chromatosome. This observation is consistent with a study showing that a nucleosome with a single linker arm is a minimal substrate for H1 binding^23^. However, the binding probability and the breaking force of GD and CTD interactions were significantly reduced (Fig. 4e,f), favoring the scenario in which CTD and GD interact with both linkers simultaneously. This was not the result of H1 dissociation from the nucleosome, since GD-DNA interaction was readily detected even after 80 cycles (>10 min) under the STO conditions.

Notably, the elimination of the distal linker, which reduced the breaking forces and binding probabilities of both CTD1 and CTD2 at the proximal linker, also reduced the rupture force of the H3-NTD contact. This might indicate that the binding of CTD to both linkers stabilizes the H3-NTD interaction. To address this, we calculated the conditional binding probability for each interaction, given that one of the other interactions is present in the same probing cycle. For WT chromatosomes, the probability of H3-NTD contact was ~2.5 fold lower than the conditional probability of an H3-NTD contact given that either a CTD1 or CTD2 interaction is present (Fig. S3b). This effect was intensified for Δ5’ chromatosomes, suggesting a tight correlation between CTD binding and H3-NTD compaction. Remarkably, the reciprocal increase in CTD binding probability by the concurrent stabilization of H3-NTD implies cooperativity between stabilization of the nucleosome and binding of the CTD.

The fact that H1 survives the repetitive unzipping (i.e., disruption of all H1-linker DNA interactions) of a nucleosome harboring a single linker DNA implies that H1 interactions with another region distinct from a linker, likely the dyad, maintains H1 bound to the nucleosome. H1 might be in a different conformation, from which it can rapidly regain its native conformation upon rezipping of the DNA. Collectively, these results indicate that the GD and the CTD bind to both linkers simultaneously, stabilizing H1 ‘on-dyad’, and that deletion of a linker DNA switches H1 to a two-contact conformation which is characterized by a decreased degree of compaction.

### Mechanical invasion into chromatosome triggers decompaction

Chromatosomes formed on the single-linker nucleosomes showed a conformation distinct from the ‘on-dyad’ one of double-linker, WT nucleosomes. We next wondered whether WT nucleosomes were able to switch their conformation following a perturbation reducing H1-DNA interactions with one of the linkers, in a manner similar to the progression of a motor protein, such as RNA polymerase, into the chromatosome. To address this, we perturbed a chromatosome by increasing the number of partial unzipping cycles from ~10 to ~80 (Fig. 5a).

**Figure 5.**
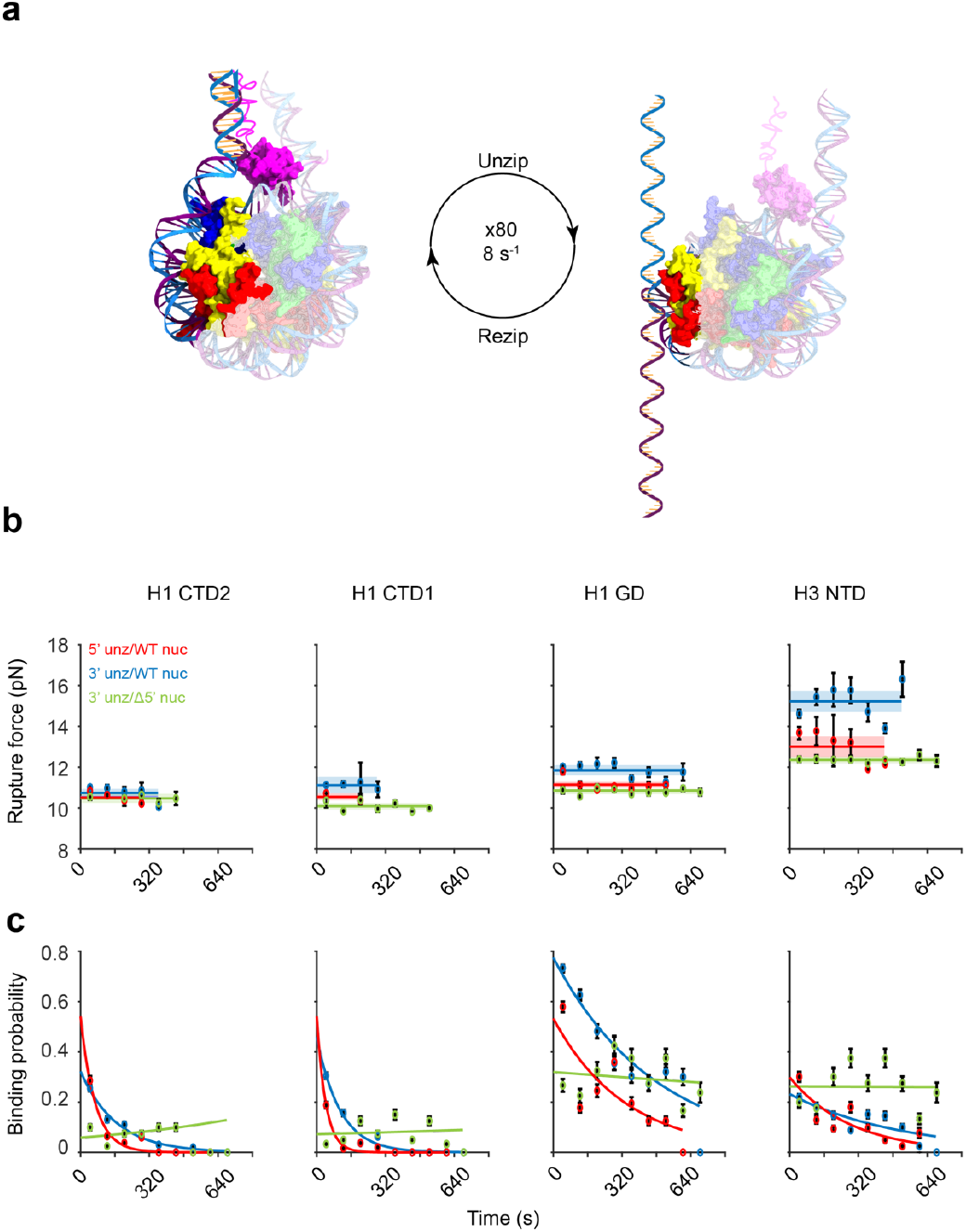
Mechanical invasion into a chromatosome triggers decompaction. a) Schematic representation of the invasion experiment. The experiment was conducted similarly to the one shown in Fig.3a but with >80 unzipping cycles. b) Rupture forces and c) binding probabilities are shown as a function of time (~720s, 80s window) for all interaction clusters of the WT and truncated chromatosomes. Data are shown as mean±s.e.m.; n_3’unz/WT nuc_ = 787, n_5’ unz/WT nuc_ = 444, n_3’ unz/Δ5’ nuc CTD_ =354, n_3’ unz/Δ5’ nuc_ =534. Exponential fits are shown to guide the eye.

Single-linker chromatosomes, for which only a two contact conformation is available, are intuitively expected to dissociate when repetitively perturbed over a long time. However, we found relatively consistent values of rupture forces (Fig. 5b) and binding probability (Fig. 5c) overtime, at all clusters of interaction, indicating that the conformation remained the same throughout the experiment. In contrast, while the mean rupture forces of all regions did not change significantly for WT nucleosomes, we observed a clear decay in binding probability as a function of time for all four regions, suggesting that H1 gradually changes its conformation. Hence, prolonged perturbation of the interactions in one linker DNA can induce a global conformational change, provided that an alternative conformation exists.

Interestingly, although the decrease in binding probability for WT chromatosomes was evident in both orientations, it was more pronounced for the perturbation from the 5’ side (Fig. 5c). Moreover, when we irreversibly unzipped the chromatosomes after the invasion from the 3’ and 5’ side, the decrease in the compaction of interactions was more substantial for the 5’ side (Fig. S4b). This suggests that the propagation of a motor protein into a chromatosome can displace H1 or its domains, leading to its decompaction, and it does so in an asymmetrical manner, with the chromatosome being more sensitive to invasion from the 5’ end.

## Discussion

In this study, we have used single-molecule optical tweezers to sequentially unzip DNA harboring a chromatosome, with the aim of measuring the position and strength of H1-DNA interactions. Our experimental system overcomes several important limitations of preceding studies: First, the usage of a laminar flow cell enables the single molecule characterization of H1-nucleosome complexes in native conditions, which otherwise rapidly disassemble. Second, the directionality and reversibility of the unzipping reaction allow measurement of the position, strength and dynamics of protein domains in the intact complex, and individual perturbation of their function by prolonged unzipping. Third, measurement of the forces associated with protein-DNA interactions provides direct assessment of the degree of compaction by H1 with high resolution and sensitivity.

Using this approach, we observed a previously unreported tightening of the H2A/H2B and H3/H4 interactions with the DNA upon H1 binding. This stabilization likely stems from the interactions of H1 at the nucleosome dyad. Two main observations support this idea: First, probing the dyad interactions using unzipping requires prior disruption of H1 interactions with the proximal linker DNA. Increased resistance to the unzipping fork at the region of proximal H2A/H2B contact suggests that H1 remains bound to the dyad, stabilizing the dimer from ‘inside-out’. The second observation is that the K69A mutation abolishes the stabilization of H2A/H2B, located ~40 bp from the center and hence not directly in contact with H1; this strongly suggests that the H1 binding site at the nucleosome center is crucial for stabilizing the whole of the nucleosome’s core. This implies that H1 binding induces a conformational change in the nucleosome dyad that propagates for at least 40 bp, most probably also affecting the linker DNA. The symmetric stabilization of both H2A/H2B dimers and entry/exit suggests that H1 binds in the on-dyad conformation. The K69A mutation leads to disruption of this symmetry, such that only half of the nucleosome is stabilized. In addition, the interaction with one of the linkers was almost abolished, indicating that most of the H1 mass is associated with one side of the nucleosome.

In contrast, the deletion of the entire CTD, which encompasses almost half of H1, only slightly destabilizes the dyad interactions and, more substantially, lessens the compaction of nucleosome entry and exit. This effect stems from the complex structure formed by CTD binding to both linker DNAs, using multiple dynamic contacts to stabilize the connections of the globular domain and the nucleosome entry and exit. The CTD interacts with both linker DNAs, making contacts ~115 (CTD1) and ~140 bp (CTD2) away from the dyad, suggesting that the absolute length of the DNA is associated with histones is ~280 bp. This is significantly longer than previously interpreted by nuclease digestion experiments^7,12,55^, and has important implications for the formation of chromatin fibers^56^ and the promotion of phase separation by linker histones^10^. The position of the proximal CTD1 contact is in agreement with an hydroxyl-radical footprinting study, which predicted that the seven amino acids in the vicinity of the CTD are responsible for the maintenance of the characteristic stem structure^17^. The previously uncharacterized CTD2 contact is located ~60 bp from the entry and exit. Its relatively fixed position, and the ability to rapidly recover following rezipping, suggest that CTD2 is ordered rather than disordered. Moreover, its distant location and dynamic nature can explain why it has not been observed in previous studies using only shorter DNA^24,26^.

The disruption of CTD1 and CTD2 contacts with DNA, either by their deletion or phosphorylation of the TPKK, changes the nature of neighboring contacts, leading to a decrease in rupture forces in GD-linker DNA contacts. This further affects the proximal H3-NTD interaction at the nucleosome entry, eventually leading to dissociation of H1. The CTD forms a similar structure with both DNA linkers, but the binding probability of these contacts at the distal linker is substantially different, suggesting that they detach more frequently. Although dynamic, these contacts play a crucial role in stabilizing the connections with the more proximal linker and in maintaining the on-dyad mode. The depletion of the distal linker by deletion or extensive unzipping, leads to a dramatic decrease in binding probability and breaking force of all interactions along with the linker DNA, resulting in destabilization of the H3-NTD contact. Strikingly, H1 does not dissociate from the nucleosome, but adopts a different conformation, making contacts with the dyad that are strong enough to survive the prolonged unzipping. In contrast, the contacts and structures formed with the linker DNA become more dynamic, leading to decreased compaction. We propose that linker depletion leads to the transition of H1 to the off-dyad mode, making transient contacts with a single DNA linker and leading to reduced nucleosome compaction.

Remarkably, our results also suggest that the cellular machinery may exploit the positional plasticity oh H1 to push it to the more dynamic and less compact ‘off-dyad’ mode, by modulating H1 contacts with the linker DNA (Fig. 6). The tendency of H1 to dissociate more rapidly from the distal side, suggests that the transition of H1 can be triggered more quickly at this location. While both H1 dissociation and downstream movement can explain the observation, it is more likely that H1 remains bound to the nucleosome, which is consistent with previous studies^56,57^. The fact that H1 survives the unzipping when bound to a single-linker nucleosome, provides and additional evidence for the resilience to DNA unzipping of an H1 contact at the dyad.

**Figure 6.**
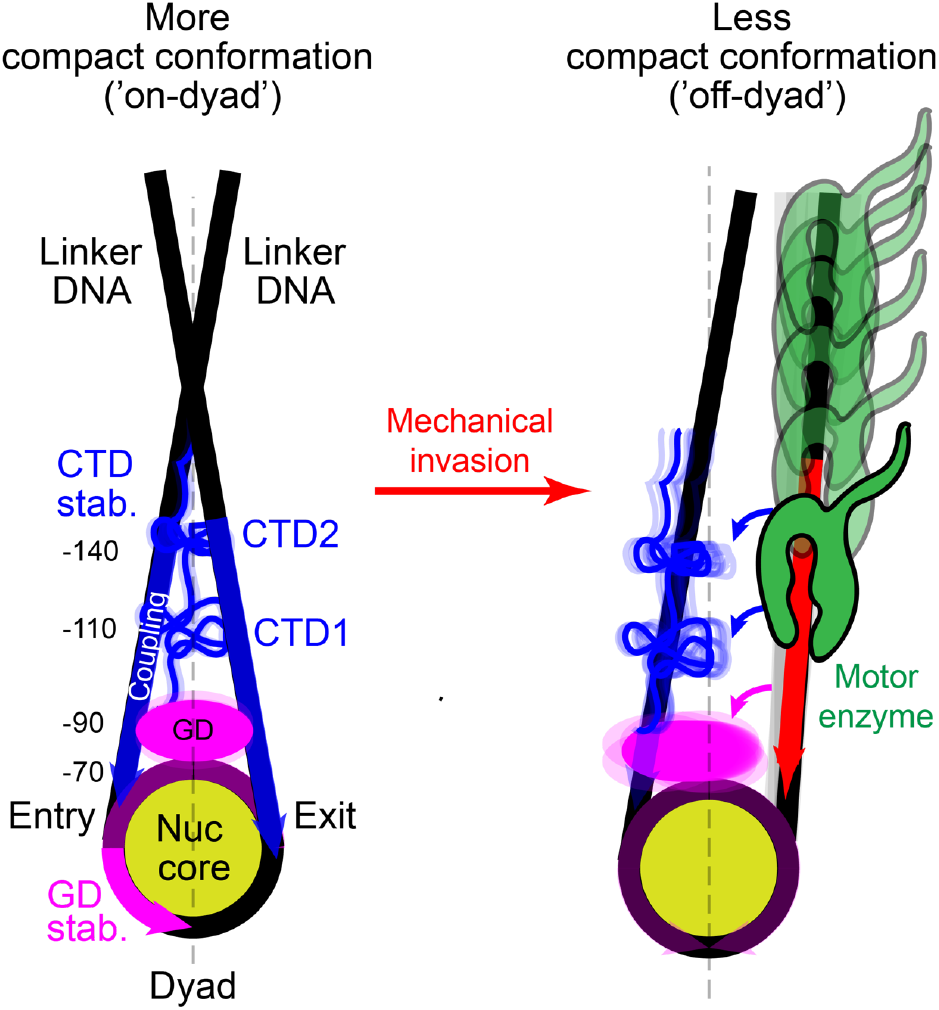
A model for dynamic compaction of a chromatosome particle. H1 binds the canonical nucleosome in an ‘on-dyad’ conformation. Binding of the GD domain to the dyad induces a conformational change leading to compaction of both H2A/H2B dimers. The CTD dynamically couples both linkers at two positions, ~±110 and ~±140 bp from the dyad, to stabilize GD-linker interactions. Mechanical invasion into the linker DNA up to the GD-DNA contact triggers H1 repositioning to the less condensed and dynamic ‘off-dyad’ conformation.

Collectively, our study suggests that a single H1 subtype can adopt both the on-dyad and off-dyad orientations, with the on-dyad being more energetically favorable^11,58^. As the off-dyad mode is associated with a less compact nucleosome, transition to this state can be fine-tuned by cellular cues to directly regulate H1 contacts at the dyad or with the linker DNA. Chromatosome orientation itself can also play a role since decompaction is triggered more easily from the distal linker.

The reduced compaction by a mimic of CTD phosphorylation suggests that some of the various H1 post-translational modifications (PTMs) that occur in dyad and linker-DNA interfaces may also affect its binding orientation^26,59–62^. For instance, modifications such as phosphorylation of the conserved Arginine 74, which is in tight contact with linker DNA or acetylation of Lysine 73, which mapped to the dyad, can lead to electrostatic repulsion with the DNA, pushing H1 to the less energetically favorable off-dyad mode. The same is true for PTMs of the core histones that can trigger decompaction by modulating interactions with the linker DNA. For example, H3K56ac that increases nucleosome unwrapping, was shown to facilitate the binding of a transcription factor to a chromatosome^40^. The fact that the binding conformation can be modified by diverse mechanisms, including PTMs, length of DNA linker and stereochemical constraints may explain the distinct binding modes observed for the same variant in different studies^16–21^.

From a broader perspective, dynamic changes in the H1 contact orientation likely affect the higher-order assembly of neighboring chromatosomes into chromatin fibers, as predicted by theoretical modeling^63^ and simulations^25^, and shown by recent cryo-EM structures ^26,27,29^. Dependence of these transitions on the modifications mentioned above may explain the absence of the canonical higher-order structures, instead of which heterogeneous ‘clutches-like’ structures are observed *in vivo*^64,65^. Moreover, the CTD of H1 was shown to promote phase separation, in a linker DNA length-dependent manner^10^. This suggests a possible role for the extended interactions we report here in controlling the effective DNA linker length, and thus regulating phase separation. Since various biological processes rely on precise control over DNA readout, the ability of a cell to alter the chromatin compaction by directly modulating the chromatosome provides a means of efficient spatiotemporal control over genetic information, which is required to orchestrate a complex physiological outcome^66^. Our work highlights how the inherent conformational plasticity of the fundamental packaging unit of chromatin may contribute to this process.

## Materials and Methods

### Histone proteins

The mH1.0 ORF was amplified from mouse cDNA using PCR with primers listed in Table S1. The product was cloned into a pColaduet-1 expression vector (71406; Merck). The H1K69A, H1T153E, and H1ΔCTD mutations were generated from WT mH1.0 in pColaduet-1 using primers listed in Table S1. All constructs were verified by sequencing. The proteins were expressed in *E. coli* Rosetta (DE3), grown for 5 h at 37 °C in LB and induced with 1 mM isopropyl-β-D-thiogalactoside at 18 °C overnight. The proteins were purified as described previously^67^ with the following modifications: The sonicated pellets were loaded into a Hydroxylapatite, Fast Flow column (391947; Millipore) pre-equilibrated with 25 mM phosphate buffer, pH 6.8, and eluted with 25 mM phosphate buffer + 1.5 M NaCl, pH 6.8. Late fractions were pooled and diluted in a 25 mM phosphate buffer to a final concentration of 50 mM NaCl. The eluates were loaded on a Sulfopropyl (SP) Sepharose column (S1799; Sigma), washed with ten volumes of 25 mM phosphate buffer + 0.5 M NaCl, pH 6.8 and eluted with 25 mM phosphate buffer + 1.5 M NaCl, pH 6.8 for WT H1, H1K69A, and H1T153E variants. The fractions containing H1ΔCTD mutant were loaded on a SP column, washed with 50 mM Tris-Cl, 2mM EDTA, 0.3 M NaCl, pH 8 and eluted using 50 mM Tris-Cl, 2mM EDTA, 0.6 M NaCl, pH 8. All fractions were concentrated using Amicon Ultra (UFC900324; Merck), snap-frozen and kept at −80°C. All buffers were supplemented with a complete EDTA-free protease inhibitor cocktail (5056489001; Roche).

Histones H1 (M2501S; NEB), H2A/H2B (M2508S; NEB), and H3.1/H4 (M2509S; NEB) from human origin were all purchased from NEB. The H2A/H2B and H3.1/H4 were mixed in 2(H2A/H2B)_2_:1(H3/H4)_4_ molar ratio to form octamers.

### Molecular construct for single-molecule experiments

For PCR-based nucleosomal DNA segments, a ~234 bp fragment of DNA containing the Widom 601 positioning sequence and a ~249 bp fragment corresponding to the −1/+223 region of the mouse *Cga* gene^43^ were amplified using standard PCR reactions with primers listed in Table S1. The constructs were digested with DraIII-HF(R3510L; NEB) or BglI (R0143L; NEB) overnight according to the manufacturer’s instructions, purified using a QIAquick PCR Purification Kit (28106; Qiagen) and mixed with recombinant histone octamers to form mono-nucleosomes under conditions reported^43^. For ligation-based nucleosomal DNA segments, oligos listed in Table S2 were purchased from IDT and phosphorylated using T4 Polynucleotide Kinase (M0201L; NEB) according to the manufacturer’s instructions. For each type of construct, oligos were mixed in a 1:1 molar ratio with their complementary sequence, as shown in Table S2. Each mix was incubated at 90°C for 5 min and annealed in 1x T4 DNA Ligase Reaction Buffer (B0202S; NEB) by gradual cooling. Each annealing reaction led to the formation of dsDNA fragment, flanked by unique sticky overhangs designed to ligate the next dsDNA fragment in a chain. The fragments were ligated using concentrated T4 DNA ligase (M0202M; NEB) and purified using a QIAquick Gel Extraction Kit (28706; Qiagen) to form constructs listed in Table S3, which were used for nucleosome reconstitution.

Two ~2000-bp DNA handles were generated as previously reported^45^, ligated to the alignment segment amplified with primers listed in Table S1 and digested with DraIII-HF. The construct was kept at −20 °C prior to usage.

Reconstituted nucleosomes were ligated to the DNA handles using T4 DNA ligase (M0202S; NEB) and 1x Rapid Ligation Buffer (C6711; Promega) in a 3:1 molar ratio, 16 h at 4°C overnight. The full construct (i.e. handles + alignment segment + nucleosome) was incubated for 15 min on ice with 0.8 μm polystyrene beads (Spherotech) coated with anti-digoxygenin. The reaction was then diluted 500-fold in H1 buffer (10 mM Tris-Cl (pH 8), 1 mM EDTA, 50 mM NaCl, 1 mM DTT, 5% v/v glycerol and 150 ng/μl BSA). Tether formation was performed inside the experimental chamber by trapping an anti-dig bead (bound by nucleosomes) in one trap, trapping a 0.9 μm streptavidin-coated polystyrene bead in the second trap, and bringing the two beads into proximity to allow binding of the biotin tag in the nucleosomal DNA to the streptavidin on the bead. The trapped nucleosome was incubated ~30 s with 5 nM of H1 in H1 buffer to form chromatosomes, after which the complex was moved to the H1-free region and unzipped (see Fig. 1a).

The correct H1:nucleosome stoichiometry (1:1) was confirmed by the following control experiments: (1) Gel shift experiments showed a single population of chromatosomes at the concentration used in single-molecule experiments. In addition, in these conditions, ~50% of nucleosomes were H1 free, indicating subsaturating conditions. (2) The alignment sequence, which was used to monitor non-specific binding or accumulation of H1 on naked DNA, showed an identical unzipping pattern in the presence or absence of H1 (Fig. 1c, Fig. S1h). (3) The magnitude of H3-NTD rupture forces, which serve as a proxy for H1 binding, remained identical when H1 concentration was reduced fivefold (Fig. S1i).

In addition, the structural integrity of H1-nucleosome complex is supported by the following observations: (1) A similar stabilization of major histone-DNA interactions was observed using the bacterially expressed mouse H1 (mH1), as with commercially available human H1 (hH1) (M2501S; NEB), sharing 96.5% identity with mH1 (Fig. S1h,i). (2) Despite their rapid dissociation, we were able to unzip a small number of chromatosomes formed in bulk and transferred to the optical tweezers; an identical stabilization of H3-NTD was detected for these chromatosomes and the chromatosomes formed *in situ* under single-molecule conditions (Fig. S1i) (3) The pattern of stabilization was similar for all H1 proteins and DNA sequences tested (i.e., 601, *Cga*, 601-AT).

### Optical Tweezers

Experiments were performed in a custom-made dual-trap optical tweezers apparatus, as previously reported^43,45,47^. Experiments were conducted using a laminar flow cell (u-Flux, Lumicks).

### Data analysis

Calibration of the setup was done by analysis of the thermal fluctuations of the trapped beads, which were sampled at 100 kHz. Experimental data were digitized at a 2500 Hz and converted into force and extension vectors using the calibration parameters. To precisely determine and subtract any residual offset in the extension of individual experiments, force-extension curves were fitted, up to 15 pN, to an extensible worm-like-chain (eWLC) model of double-stranded DNA with persistence length 45 nm, contour length per base pair 0.34 nm/bp and stretch modulus 1000 pN. Then, to calculate the number of unzipped base pairs we divided the extension by twice the contour length of a ssDNA nucleotide, calculated with a WLC model with persistence length 1 nm and contour length per nucleotide 0.64 nm/nt. The 248 bp naked DNA alignment segment was used to perform a correlation-based alignment of all traces in an experiment. The ‘distance from dyad’ in force-position curves was calculated by subtracting the known position of the nucleosome’s center at each construct: 360 bp for a 601 DNA, and 486 bp for 601-AT.

In full, irreversible unzipping experiments (Fig. 1, Fig. 2, Fig. S1a,b,c,e,h, Fig.S4) the steerable trap was continuously moved at 280 nm/s to stretch the tethered construct, until the nucleosome fully disassembled. In the repetitive/reversible unzipping experiments (Fig. 3,4,5, Fig. S1i, S2, S3), the steerable trap was moved only until the fork reaches the point where the nucleosome’s proximal H2A/H2B is identified (typically ~19-22 pN). At this point, the steering direction is reversed, thus relaxing the force and allowing the DNA to reanneal. Each tethered complex was first reversibly unzipped 10 times in the absence of H1 (control experiment) and then reversibly unzipped 10-80 times in the presence of H1 (experimental traces), with a cycle time of 8 s, unless specified. Data were low-pass filtered for further analysis with a zero phase Butterworth filter, with a bandwidth of 40 and 150 Hz for the irreversible and reversible unzipping experiments, respectively.

Dwell time histograms (Fig. 1f,g; Fig. 2b,c) were calculated by counting the number of data points in 1 bp bins, and dividing by the sampling rate (2500 Hz). Averaged force-position traces (Fig. 2d,e; Fig. S4b) were calculated by resampling (i.e. interpolating) the force vectors of different traces into a common uniformly-spaced 1 bp array, and averaging the interpolated forces.

To calculate the mean breaking force in a specific region, we generated an “interaction vector” for each trace, where an interaction is defined as an increase in force at constant position, followed by a force drop (a break). Dwell time histograms were used to detect clusters of interactions corresponding to specific histone-DNA interactions (i.e. H3-NTD, H2A/H2B, H3/H4 for Fig. 1 and Fig. 2) and determine their boundaries. The interaction with the highest breaking force inside the boundaries of each region defines the region’s breaking force, which is then averaged over the ensemble.

Given the clear separation of rupture forces in experiments with or without H1 in all clusters of interactions (Fig. S5a,b), we identified “bound” events as interactions with higher breaking force than that of the same region in the control experiments, plus twice its standard deviation. Applying the same criteria for the data obtained without H1 resulted in detection of less than 1.5 % of the events detected with H1. The binding probability is defined as the number of bound events detected at a specific cycle out of the total number of experiments and is shown only for data containing at least 5 experiments. Mean forces were calculated by averaging the breaking force of all the detected bound events in a certain region. Only data calculated from more than 2 data points (i.e. 2 bound events) are shown in Fig. 3c and Fig. S2d. For the longer invasion experiments, the data were calculated over ten-cycle windows in order to increase the data points at each window. The mean breaking force is presented only if more than two data points were identified as being bound at each window, and the probability is presented only if at least 10 experiments exist.

## Author Contributions and Notes

S.R. and Y.G performed the experiments. S.R. and E.G. prepared experimental materials. S.R. and H.K. analyzed the data. S.R. and A.K. wrote the paper. P.M and A.K. supervised the research.

The authors declare no conflict of interest.

## Acknowledgments

We acknowledge support from the Israel Science Foundation (Grants 1782/17 to A.K. and 1850/17 to PM).

## Notes

### Competing Interest Statement

The authors have declared no competing interest.

