## Supplementary Information for "Extended and dynamic linker histone-DNA interactions control chromatosome compaction" (from another version of this preprint: version 1, 2020-10-10)

### SUPPLEMENTARY FIGURES

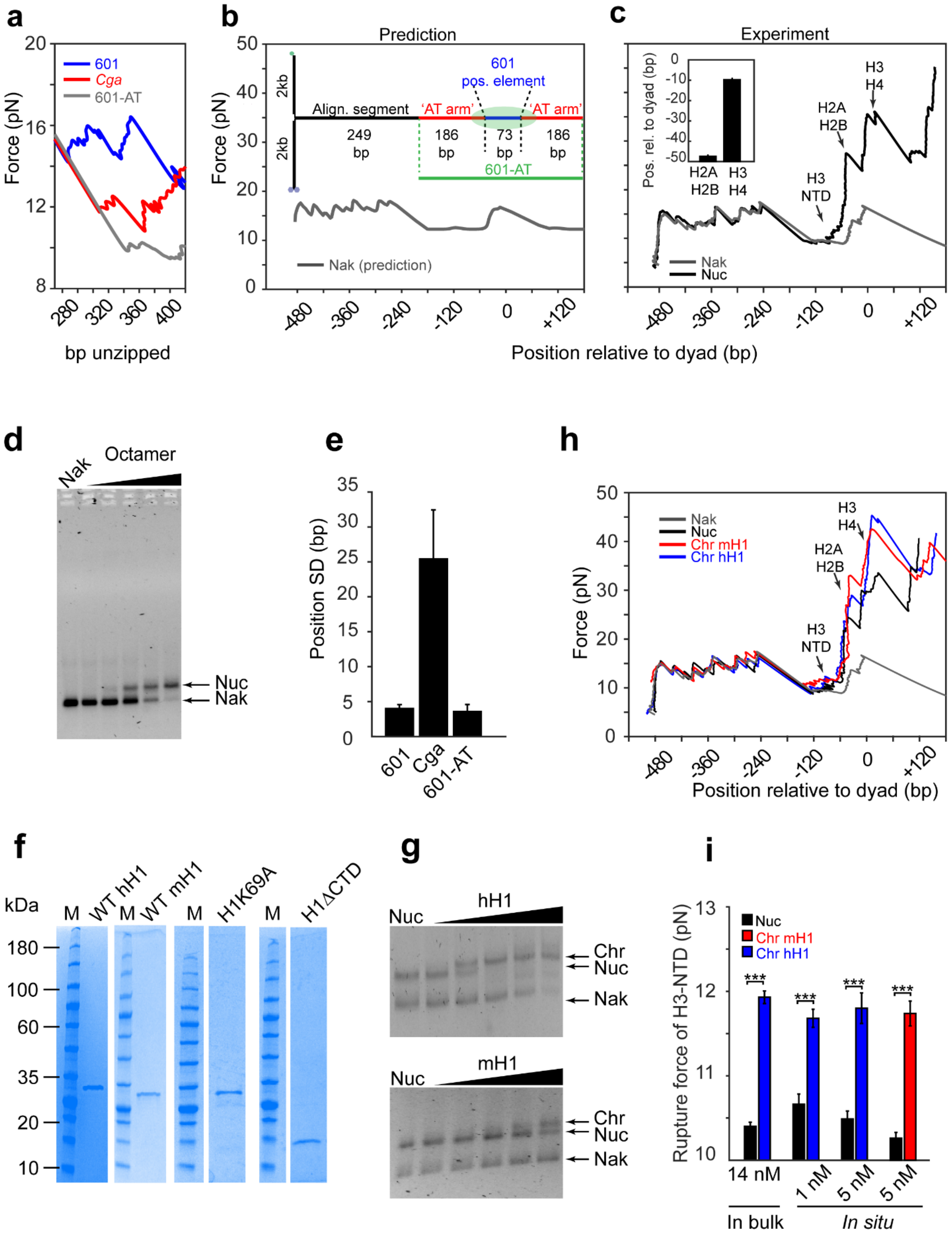

**Figure S1. The integrity of nucleosomes and chromatosomes formed on the 601-AT construct. a)** Unzipping traces for naked DNA corresponding to the 601 (blue), *Cga* (red), and 601-AT (grey) constructs. **b)** Simulation for the unzipping reaction of the construct composed of the ~249 bp ‘alignment segment’ ligated to ~471 bp 601-AT DNA. The construct is connected to ~4kb DNA handles bound to trapped beads using terminal modifications. **c)** Representative unzipping curves for naked (grey) or nucleosomal 601-AT DNA (black). Note a conserved ~35-40 bp distance between the H3/H4 and H2A/H2B region in the population of 601-AT nucleosomes. Data are shown as mean±s.e.m., $n_{601\text{-}AT}$=53. **d)** Typical nucleosome reconstitution using 601-AT DNA shows a single population of nucleosomes. DNA was titrated with increasing amounts of histone octamers for each reconstitution reaction. All reconstitutions were performed under the same conditions, and 1/4 of each reconstitution reaction was run on a 1% agarose gel post-stained with EtBr. ‘Nak’ stands for naked DNA – with no histones. **e)** Positional dispersion of the H2A/H2B interaction in 601, *Cga*, and 601-AT nucleosomes. Data are shown as s.d.±s.e., *n*=13, 14, and 53, respectively. **f)** H1 proteins used in the study. WT mouse linker histone subtype H1.0 (mH1) and mutant proteins were expressed in bacteria and purified using hydroxyapatite (HA) and sulfopropyl (SP) columns under native conditions as reported[49] (see Methods). Commercially available human H1.0 (hH1) was purchased from New England Biolabs (M2501S; NEB) and is shown for reference. The preps were resolved on separate 2-20% gradient SDS-PAGE (456-1094; BioRad). **g)** Formation of chromatosomes on 601-AT DNA. Reconstituted nucleosomes were incubated 30 min on ice in H1 buffer (see Methods) with increasing amounts of hH1 (top panel) or mH1 (bottom panel) and resolved on a 1% agarose gel post-stained with EtBr. **h)** Representative unzipping curves for naked (grey), nucleosomal 601-AT DNA (black), and nucleosomes incubated *in situ* with hH1 (blue) and mH1 (red). **i)** Similar stabilization of the H3-NTD region for chromatosomes formed in bulk (as shown in Supp.Fig.1g) and measured at single-molecule conditions (for the rare cases where a chromatosome could be detected before dissociation) and chromatosomes formed *in situ,* using different concentrations and paralogs of H1. The chromatosomes were probed using a reversible unzipping assay (as described in Fig.3a). Data are shown as mean±s.e.m.; $n_{bulk\_nuc}$=114, $n_{bulk\_hH1}$=209, $n_{in\_situ\_nuc\_1nM}$=62, $n_{in\_situ\_hH1\_1nM}$=110, $n_{in\_situ\_nuc\_5nM}$=50, $n_{in\_situ\_hH1\_5nM}$=69, $n_{in\_situ\_nuc\_5nM}$=187, $n_{in\_situ\_mH1\_5nM}$=242. ***$P$<0.001, two-sample Student’s *t*-test.

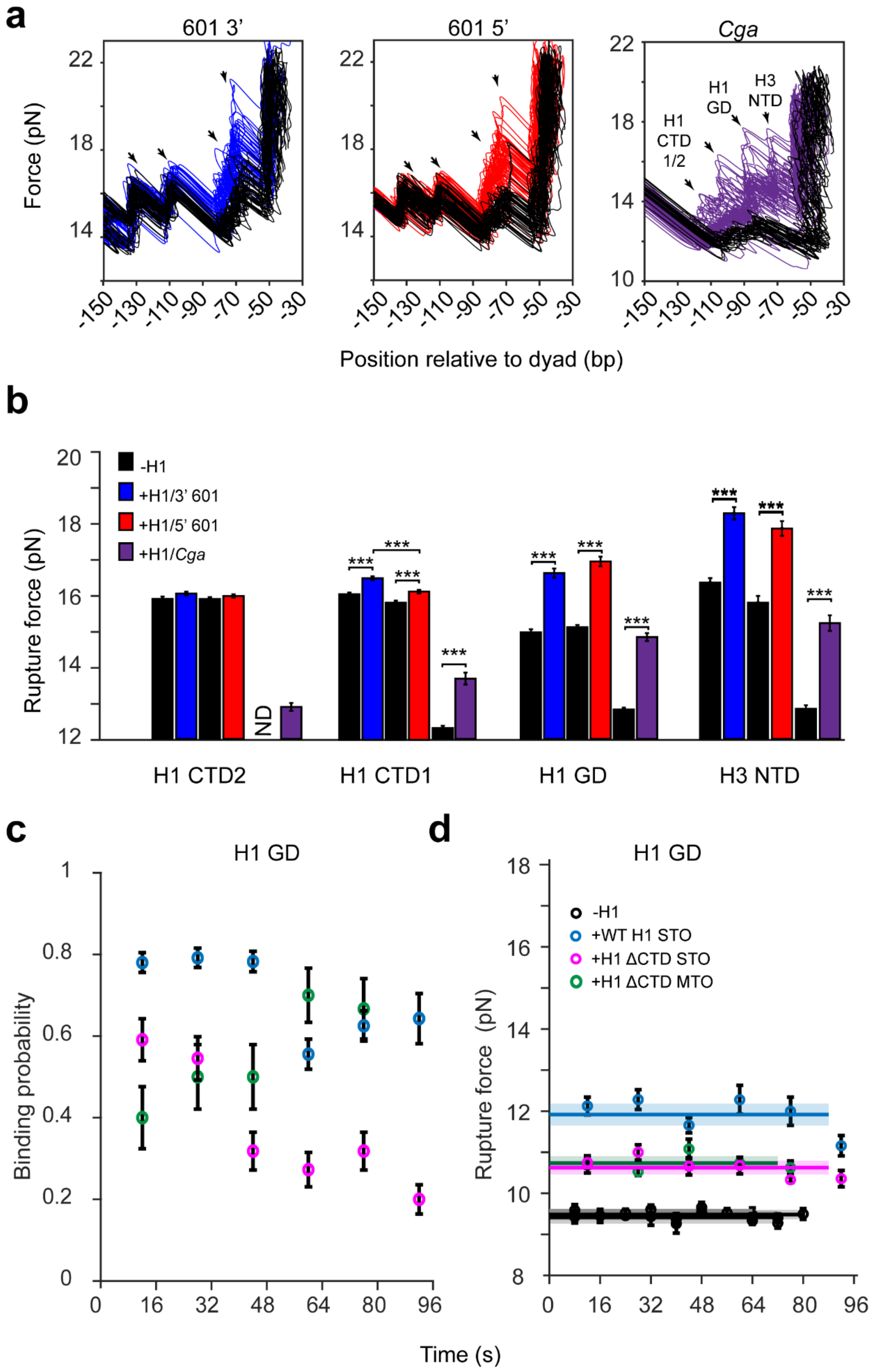
a
601 3'
601 5'
Cga
Force (pN)
H1 CTD 1/2
H1 GD
H3 NTD
Position relative to dyad (bp)
b
-H1
+H1/3' 601
+H1/5' 601
+H1/Cga
Rupture force (pN)
ND
H1 CTD2
H1 CTD1
H1 GD
H3 NTD
c
H1 GD
Binding probability
d
H1 GD
-H1
+WT H1 STO
+H1 ΔCTD STO
+H1 ΔCTD MTO
Rupture force (pN)
Time (s)

**Figure S2. A similar trend of linker DNA-H1 interactions for 601 and *Cga* chromatosomes**. **a)** Repetitive unzipping force-extension curves of chromatosomes formed on 601 (unzipped from 3' end–blue or 5' end–red) or *Cga* nucleosomes (purple). Traces of nucleosomes before the addition of H1 are shown in black. The clusters of detected interactions are highlighted with arrows. **b)** The contacts detected in each unzipping cycle were analyzed and presented as mean rupture forces. Data are shown as mean±s.e.m.; $n_{5'601}$= 53, $n_{5'601+H1}$= 60, $n_{3'601}$=37, $n_{3'601+H1}$=47, $n_{Cga\_nuc}$=27, $n_{Cga+H1}$=59. **c)** Binding probabilities and **d)** rupture forces of the GD cluster as a function of time. WT chromatosomes (blue) were probed under STO conditions. H1△CTD chromatosomes were analyzed under STO conditions (purple) or MTO conditions (green). Rupture forces for WT STO and H1△CTD MTO are presented for comparison. Data are shown as mean±s.e.m.; $n_{nuc\_ctrl}$=252, $n_{nuc+WTH1\_MTO}$=307, $n_{nuc+H1\triangle CTD\_STO}$=41, $n_{nuc\_ctrl\_\triangle CTD\_MTO}$=55, $n_{nuc+H1\triangle CTD\_MTO}$=51. ****P*<0.001, two-sample Student's *t*-test.

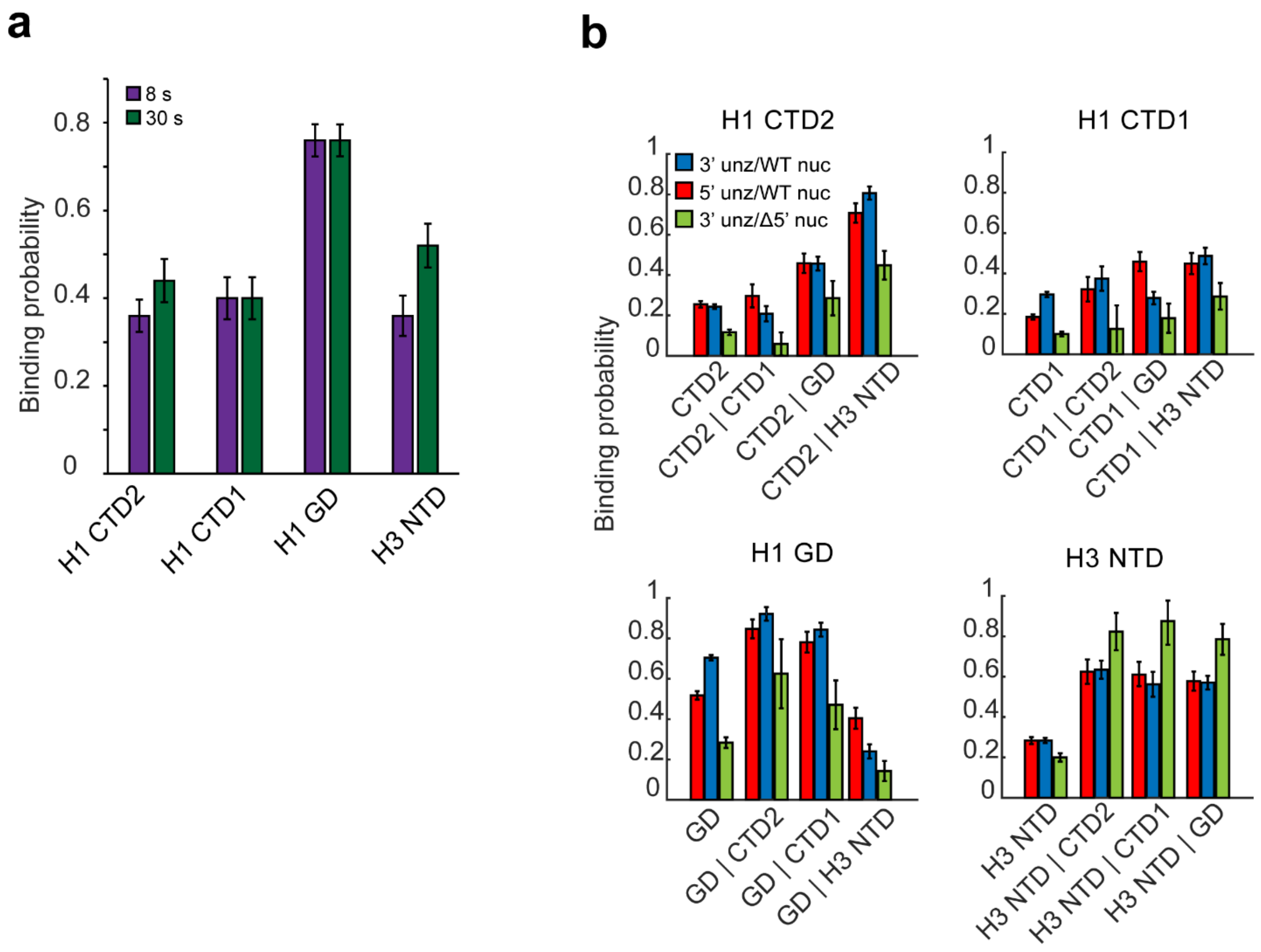


**Figure S3. Determination of the binding probability of histone-DNA interactions on linker DNA. a)** Binding probability in each region of interaction for a population of WT chromatosomes probed after a typical incubation time of 30 s (green) or 8 s (purple), in the 3' direction. Data are shown as mean±s.e.m.; $n_{8s}$ =25, $n_{30s}$ =25. **b)** Binding probabilities for all regions, both for the complete data set as presented in Fig.4f as well as for conditional subsets (*r*|*s* indicates probability of binding of interaction *r* given that interaction *s* is bound too). The data is analyzed and presented as in Fig. 4f.

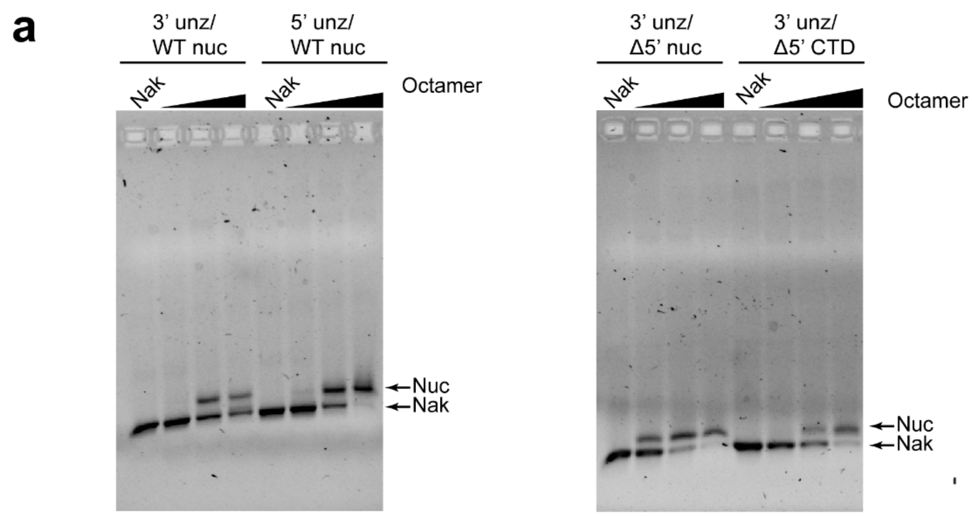


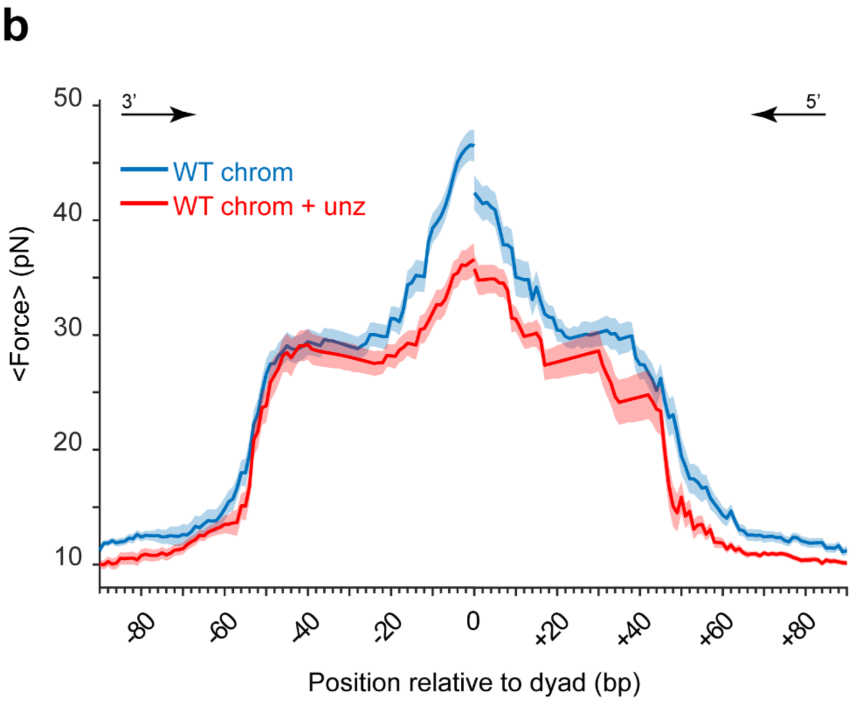


**Figure S4. Mechanical invasion into chromatosome triggers decompaction. a)** Typical reconstitution for the nucleosomal constructs used in Fig. 4 and Fig. 5. DNA was titrated with increasing amounts of histone octamers for each reconstitution reaction. 1/4 of each reconstitution reaction was run on a 1% agarose gel post-stained with EtBr. 'Nak' stands for naked DNA – with no histones. **b)** Mean force-position curves for chromatosomes probed irreversibly following ≥10 cycles of reversible unzipping, either from 5' or 3' direction (red). Mean rupture force-position curves for unperturbed chromatosomes are shown as a reference (blue). Data are shown as mean±s.e.m.; $n_{WT3'chrom}$ =33, $n_{WT3'chrom+unz}$ =20, $n_{WT5'chrom}$ =14, $n_{WT5'chrom+unz}$ =19.

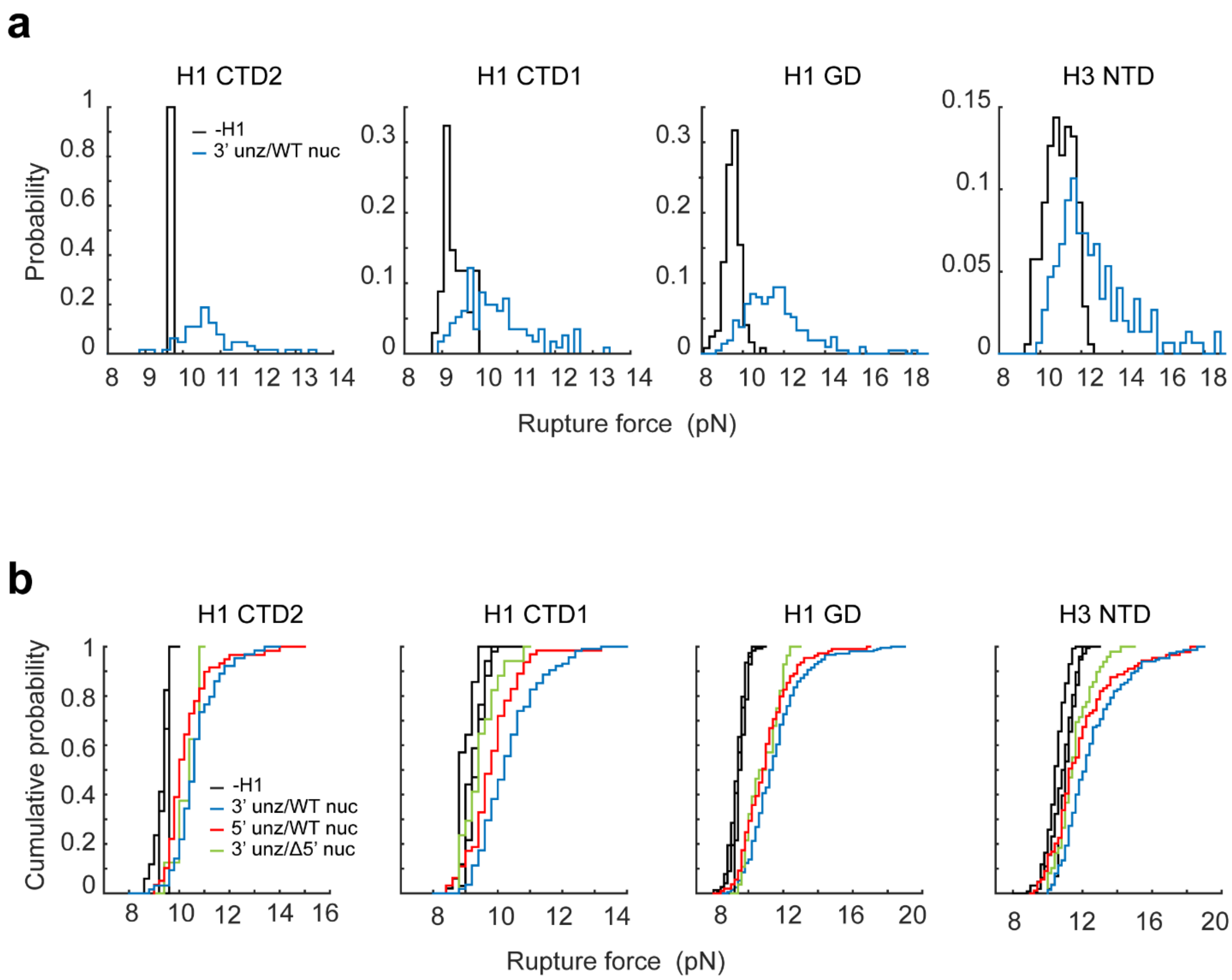


**Figure S5. Increased rupture forces of protein-linker DNA interactions upon the addition of H1.** **(a)** Probability and **(b)** cumultive probability of breaking forces for all intereactions detected in the linker DNA regions in the absence (black) and presence of H1 (blue: 3' unz/WT nuc, red: unz/WT nuc, green: 3' unz/ ∆5' nuc).

### SUPPLEMENTARY TABLES

**Table S1:** Primers used for PCR amplification of DNA sequences

| Name | DNA sequence (5'→3') |
|---|---|
| 601 bgl1 F | ATGGCCTTGCCGGCACTACGTAATATTGGCCAGCTAGGACGCCCCCAAATG TATATATCTGACACGTGCCTGGAGACTA |
| 601 dra3 R | ATCACTGCGTGCGGCACTACGTAATATTGGCCAGCTAGGACGCCCCCAAGA ATCCCGGTGCCGAGGC |
| Cga -1 bgl1 F | ATGGCCTTGCCGGCGCAGGCACTGAAAAATCCAG |
| Cga + 223 dra3 R | ATCACTGCGTGCAGCCATGAGCACCATAAAG |
| Alignment CAC DraIII F | ATCACCACGTGGAATTCGATATCCCCGAGA |
| Alignment 834 R DraIII | ATCACTGCGTGGCACGGCCGATGGACCCTATAC |
| 2K BioTg F | /5'Biotin-TEG/GATCTCCAGCCAGGAACTATTGA |
| 2K bio Nt.BvCI R | GTGTCAGCTTGCCCCTCAGCGATGACCTCAGCATTTTTCGACCTGCTCTTC AGCA |
| 2K DoubleDig2 Bgl1F | ATGGCCTAGACGGCGAGCCTGGGTTTATAAGGGGAGCGGTGA |
| 2K Dig Nb.BbVCI R | TCAGCTTGCCCCTCAGCGATGACCTCAGCAAGGACCAGCGTTTTGTTGAAA |
| Oligo 5' Dig AGA | /5'Digoxigenin(NHS ester/AGGTGCCGGGACCACCCTGTAGA |
| Oligo 3' Dig/ 5' phos | /5'Phosphate/ACAGGGTGGTCCCGGCACCT/3'Digoxigenin (NHS ester/ |
| mH10 WT NdeI F | CATATGACCGAGAACTCCACCTCCGC |
| mH10 WT XhoI R | CTCGAGTTATCACTTCTTCTTGCTGGCCCTCT |
| mH10 K69A R | TCGGCGTTCTCACCCACC |
| mH10 K69A F | CTCCCAGATCGCGTTGTCCATCAAGCGCC |
| mH10 Δ CTD 97 R | CTTGGCCAGCCTGAAGGA |
| mH10 Δ CTD 97 F | TGATAACTCGAGTCTGGTAAAGAAAC |
| mH10 T153E F | GCCGGCTGCCgagCCCAAGAAAG |
| mH10 T153E R | TTCTTCTTGGCCTTCTTGAC |

| Name | DNA sequence (5'→3') | 3'unz/ WT nuc | 5'unz/ WT nuc | 3'unz/ Δ5'nuc |
|---|---|---|---|---|
| O1 F OT dra3 | pGTGCGGCATGAAGACTATAAATTAATATTAATATTTATATATAA AATTATTTAATATAATTATAAATATTTT | + | + | - |
| O1 R dra3 | pTATTTATAATTATATTAAATAATTTTATATATAAATATTAATAT TAATTTATAGTCTTCATGCCGCACGAC | + | - | - |
| O1 R OT dra3 | pTATTTATAATTATATTAAATAATTTTATATATAAATATTAATAT TAATTTATAGTCTTCATGCCGCACGCA | - | + | - |
| O2 F | pTAAATTATATTTTTATAAATTATTATAAAATTTTTAATTTTAAT TATTACATTATTTTAATATTAAATATAAAAATAA | ++ | ++ | + |
| O2 R | pTTTTATATTTAATATTAAAATAATGTAATAATTAAAATTAAAAA TTTTATAATAATTTATAAAAATATAATTTAAAAA | ++ | ++ | + |
| O3 F | pAAATTATTTAATTAAATATTTATAAATTATTTTATAATATTATA TATATGGAGTAATCCCCTTGGCGGTTAAAACGCGGG | + | + | + |
| O3 R | pCGTTTTAACCGCCAAGGGGATTACTCCATATATATAATATTATA AAATAATTTATAAATATTTAATTAAATAATTTTTAT | + | + | + |
| Rev 3F | pGGACAGCGCGTACGTGCGTTTAAGCGGTGCTAGAG | + | + | + |
| Rev 3R | pTAGCACCGCTTAAACGCACGTACGCGCTGTCCCCCG | + | + | + |
| Rev 4F | pCTTGCTAATATATATAATATTATAAAATAATTTATAAATATTTA ATTAAATAATTTTTAT | + | + | + |
| Rev 4R | pAAATTATTTAATTAAATATTTATAAATTATTTTATAATATTATA TATATTAGCAAGCTC | + | + | + |
| Rev 6 R OT bgl1 | pCGGCGGCATGAAGACTATAAATTAATATTAATATTTATATATAA AATTATTTAATATAATTATAAATATTTT | + | + | + |
| Rev 6 F bgl1 | pTATTTATAATTATATTAAATAATTTTATATATAAATATTAATAT TAATTTATAG TCTTCATGCCGCCGGAC | - | + | - |
| Rev 6 F OT bgl1 | pTATTTATAATTATATTAAATAATTTTATATATAAATATTAATAT TAATTTATAGTCTTCATGCCGCCGGCA | + | - | - |
| Δ 5 nuc F | TTATTACATTATTTTAATATTAAATATAAAAATAA | - | - | + |
| Δ 5 nuc R | pTTTTATATTTAATATTAAAATAATGTAATAA | - | - | + |

**Table S2:** Oligos used for the formation of 601-AT based constructs

- Not used, + used once, ++ used twice

p – phosphorylated

**Table S3:** PCR and oligo-ligation products used for the formation of unzipping constructs.

| Name | DNA sequence (5'→3') |
|---|---|
| Alignment sequence<br>(248 bp) | GGTCATCGCTGAGGGGCAAGCTGACACGTGGAATTCGATATCCCCGAGAAG<br>GTCGCTGTTCAATACATGCACAGGATGTATATATCTGACACGTGCCTGGAG<br>ACTAGGGAGTAATCCCCTTGGCGGTTAAAACGCGGGGGACAGCGCGTACGT<br>GCGTTTAAGCGGTGCTAGAGCTTGCTACGACCAATTGAGCGGCCTCGGCAC<br>CGGGATTCTCCAGGGCGGCCGCGTATAGGGTCCATCGGCCGTGC |
| 601 DNA<br>(234 bp) | ATGGCCTTGCCGGCACTACGTAATATTGGCCAGCTAGGACGCCCCCAAATG<br>TATATATCTGACACGTGCCTGGAGACTAGGGAGTAATCCCCTTGGCGGTTA<br>AAACGCGGGGGACAGCGCGTACGTGCGTTTAAGCGGTGCTAGAGCTTGCTA<br>CGACCAATTGAGCGGCCTCGGCACCGGGATTCTTGGGGGCGTCCTAGCTGG<br>CCAATATTACGTAGTGCCGCACGCAGTGAT |
| Cga DNA<br>(249 bp) | ATCACTGCGTGCAGCCATGAGCACCATAAAGAAAATTAATGAATGCCTAAT<br>TATTTTCCTACCAACATTTATGCAGAAAATTCTTTCCTGTGGTCACAAATA<br>TTTTACTCTTTAATTAAATCCCCCCAAATGATTTCATCATATTACCTACCT<br>CTGGACTTTTTGAGGGATGTGTTCTGGGCAGGTGATTGTCGATCTGAGGGG<br>AACAATGTCTCTGGATTTTTCAGTGCCTGCAGCCGGACAGGCCA |
| 601-AT – 3'unz/WT nuc<br>601-AT – 5'unz/WT nuc<br>(471 bp) | GTGCGGCATGAAGACTATAAATTAATATTAATATTTATATATAAAATTATT<br>TAATATAATTATAAATATTTTTAAATTATATTTTTATAAATTATTATAAAA<br>TTTTTAATTTTAATTATTACATTATTTTAATATTAAATATAAAAATAAAAA<br>TTATTTAATTAAATATTTATAAATTATTTTATAATATTATATATATGGAGT<br>AATCCCCTTGGCGGTTAAAACGCGGGGGACAGCGCGTACGTGCGTTTAAGC<br>GGTGCTAGAGCTTGCTAATATATATAATATTATAAAATAATTTATAAATAT<br>TTAATTAAATAATTTTTATTTTTATATTTAATATTAAAATAATGTAATAAT<br>TAAAATTAAAAATTTTATAATAATTTATAAAAATATAATTTAAAAATATTT<br>ATAATTATATTAAATAATTTTATATATAAATATTAATATTAATTTATAGTC<br>TTCATGCCGCCG |
| 601-AT – 3'unz/ Δ5'nuc<br>CTD<br>(321 bp) | CGGCGGCATGAAGACTATAAATTAATATTAATATTTATATATAAAATTATT<br>TAATATAATTATAAATATTTTTAAATTATATTTTTATAAATTATTATAAAA<br>TTTTTAATTTTAATTATTACATTATTTTAATATTAAATATAAAAATAAAAA<br>TTATTTAATTAAATATTTATAAATTATTTTATAATATTATATATATTAGCA<br>AGCTCTAGCACCGCTTAAACGCACGTACGCGCTGTCCCCCGCGTTTTAACC<br>GCCAAGGGGATTACTCCATATATATAATATTATAAAATAATTTATAAATAT<br>TTAATTAAATAATTT |
| 601-AT – 3'unz/ Δ5'nuc<br>CTD<br>(311 bp) | CGGCGGCATGAAGACTATAAATTAATATTAATATTTATATATAAAATTATT<br>TAATATAATTATAAATATTTTTAAATTATATTTTTATAAATTATTATAAAA<br>TTTTTAATTTTAATTATTACATTATTTTAATATTAAATATAAAAATAAAAA<br>TTATTTAATTAAATATTTATAAATTATTTTATAATATTATATATATTAGCA<br>AGCTCTAGCACCGCTTAAACGCACGTACGCGCTGTCCCCCGCGTTTTAACC<br>GCCAAGGGGATTACTCCATATATATAATATTATAAAATAATTTATAAATAT<br>TTAAT |

Cyan – 73 bp core of the 601 positioning sequence

Green – nucleosomal DNA surrounding the 73 bp core

Yellow - linker DNA

### **Table S4:** Calculated P-values

| **Figure 1d - Rupture forces** | Student's *t*-test | | |
|---|---|---|---|
| 601 | H3 NTD | H2A H2B | H3 H4 |
| Nuc vs Nuc +H1 | 0.00027 | 6.00E-05 | 0.1 |
| **Figure 1e - Rupture forces** | Student's *t*-test | | |
| *Cga* | H3 NTD | H2A H2B | H3 H4 |
| Nuc vs Nuc +H1 | 0.012 | 0.037 | 0.3 |

| **Figure 2 - Rupture forces** | Student's *t*-test | | | | | |
|---|---|---|---|---|---|---|
| | 601-AT unzipped from 3' direction | | | 601-AT unzipped from 5' direction | | |
| | H3 NTD | H2A H2B | H3 H4 | H3 NTD | H2A H2B | H3 H4 |
| Nuc vs Nuc+WT H1 | 1.00E-05 | 8.00E-06 | 0.00061 | 5.00E-06 | 0.01 | 0.01 |
| Nuc vs Nuc+H1K69A | 0.002 | 0.3 | 0.3 | 2.00E-05 | 0.001 | 0.03 |
| Nuc vs Nuc+H1ΔCTD | 0.00013 | 9.00E-05 | 0.4 | 0.0009 | 0.1 | 0.1 |
| Nuc+WT H1 vs Nuc+H1K69A | 0.016 | 0.003 | 0.04 | 0.3 | 0.3 | 0.9 |
| Nuc+WT H1 vs Nuc+H1ΔCTD | 0.044 | 0.8 | 0.042 | 0.05 | 0.2 | 0.4 |
| Nuc+H1K69A vs Nuc+H1ΔCTD | 0.7 | 0.0039 | 0.8 | 0.2 | 0.04 | 0.4 |

| **Figure 3d - Rupture forces** | Student's *t*-test | | | |
|---|---|---|---|---|
| | H1 CTD2 | H1 CTD1 | H1 GD | H3 NTD |
| Nuc vs Nuc+WT H1 | 0.02 | 1.00E-15 | 2.00E-56 | 2.00E-56 |
| Nuc vs Nuc+H1T153E | 0.054 | 2.00E-13 | 1.00E-42 | 1.00E-42 |
| Nuc vs Nuc+H1ΔCTD | ND ΔCTD | 0.0018 | 6.00E-34 | 6.00E-34 |
| Nuc+WT H1 vs Nuc+H1T153E | 0.01 | 0.5 | 0.0054 | 0.0015 |
| Nuc+WT H1 vs Nuc+H1ΔCTD | ND ΔCTD | 0.0009 | 5.00E-05 | 0.0011 |
| Nuc+H1T153E vs Nuc+H1ΔCTD | ND ΔCTD | 0.0013 | 0.021 | 0.1 |

| **Figure 4e- Rupture forces** | Student's *t*-test | | | |
|---|---|---|---|---|
| | H1 CTD2 | H1 CTD1 | H1 GD | H3 NTD |
| 3' unz/WT nuc-H1 vs 3' unz/WT nuc +H1 | ND ctrl | 5.00E-10 | 1.00E-38 | 5.00E-25 |
| 5' unz/WT nuc-H1 vs 5' unz/WT nuc +H1 | 4.00E-05 | 1.00E-05 | 1.00E-19 | 1.00E-13 |
| 3' unz/Δ5' nuc-H1 vs 3' unz/Δ5' nuc +H1 | 0.017 | 0.011 | 3.00E-05 | 0.0076 |
| 3' unz/WT nuc +H1 vs 5' unz/WT nuc +H1 | 0.014 | 0.00026 | 0.0028 | 0.01 |
| 3' unz/WT nuc +H1 vs 3' unz/Δ5' nuc +H1 | 0.3 | 0.00014 | 0.025 | 0.00022 |
| '5' unz/WT nuc +H1 vs 3' unz/Δ5' nuc +H1 | 0.7 | 0.055 | 0.5 | 0.2 |

| **Figure 4f - Binding probabilities** | *χ2* test | | | |
|---|---|---|---|---|
| | H1 CTD2 | H1 CTD1 | H1 GD | H3 NTD |
| 3' unz/WT nuc-H1 vs 3' unz/WT nuc +H1 | ND ctrl | 1.00E-16 | 9.00E-49 | 2.00E-15 |
| 5' unz/WT nuc-H1 vs 5' unz/WT nuc +H1 | 2.00E-10 | 6.00E-07 | 8.00E-22 | 5.00E-12 |
| 3' unz/Δ5' nuc-H1 vs 3' unz/Δ5' nuc +H1 | 0.0064 | 0.011 | 8.00E-06 | 4.00E-06 |
| 3' unz/WT nuc +H1 vs 5' unz/WT nuc +H1 | 0.8 | 0.015 | 0.00023 | 0.9 |
| 3' unz/WT nuc +H1 vs 3' unz/Δ5' nuc +H1 | 0.032 | 0.0018 | 1.00E-09 | 0.1 |
| 5' unz/WT nuc +H1 vs 3' unz/Δ5' nuc +H1" | 0.028 | 0.1 | 0.0022 | 0.2 |

| **Figure S1i - Rupture forces** | Student's *t*-test | | |
|---|---|---|---|
| | bulk/in situ | human/mouse | H3 NTD |
| Nuc vs Nuc +H1 14 nM | bulk | human | 4.00E-50 |
| Nuc vs Nuc +H1 1 nM | in situ | human | 2.00E-10 |
| Nuc vs Nuc +H1 5 nM | in situ | human | 1.00E-07 |
| Nuc vs Nuc +H1 5 nM | in situ | mouse | 5.00E-25 |

| **Figure S2b - Rupture forces** | Student's *t*-test | | | |
|---|---|---|---|---|
| | H1 CTD2 | H1 CTD1 | H1 GD | H3 NTD |
| 3' 601 Nuc vs 3' 601 Nuc +H1 | 0.2 | 0.0001 | 9.00E-18 | 5.00E-09 |
| 5' 601 Nuc vs 5' 601 Nuc + H1 | 0.083 | 1.00E-07 | 1.00E-09 | 4.00E-13 |
| 3' 601 Nuc +H1 vs 5' 601 Nuc + H1 | 0.3 | 2.00E-06 | 0.1 | 0.1 |
| Cga Nuc vs Cga Nuc +H1 | ND ctrl | 5.00E-08 | 5.00E-14 | 0.0014 |

| **Figure S3a - Binding probabilities** | $\chi 2$ test | | | |
|---|---|---|---|---|
| | H1 CTD2 | H1 CTD1 | H1 GD | H3 NTD |
| 8 s vs. 30s | 0.5 | 1.0 | 1.0 | 0.2 |
